# Corticosterone drives behavioral inflexibility via plasticity-related gene expression in the dorsal striatum

**DOI:** 10.1101/2025.07.24.666631

**Authors:** Michael D. Murphy, Keegan S. Krick, Shuo Zhang, Elizabeth A. Heller

**Affiliations:** Department of Systems Pharmacology and Translational Therapeutics, University of Pennsylvania, Philadelphia, PA, 19104, USA; Penn Epigenetics Institute, Perelman School of Medicine, University of Pennsylvania, Philadelphia, PA, 19104, USA

**Keywords:** dorsal striatum, glucocorticoid receptor, epigenetics, sex differences

## Abstract

Behavioral flexibility allows organisms to modify actions based on new information, such as shifts in reward value or availability, and is promoted by the dorsomedial striatum (DMS). In contrast, behavioral inflexibility provides efficiency and automaticity in familiar contexts, and is promoted by the dorsolateral striatum (DLS). Importantly, chronic elevation of the primary stress hormone, corticosterone (CORT) in rodents or cortisol in humans, impairs behavioral flexibility through dendritic atrophy in the DMS, and promotes inflexible behavioral response strategies through dendritic outgrowth in the DLS. However, understanding of the molecular mechanisms that underlie the structural changes promoting behavioral inflexibility is lacking. We used a food-motivated operant task in male and female mice to define synaptic plasticity gene regulation supporting a decreased DMS activity and increased DLS activity in the shift to inflexible behavior with CORT. We discovered that CORT-accelerated loss of behavioral flexibility is accompanied by decreased DMS- and increased DLS-specific synaptic plasticity gene expression, and that distinct genes are either differentially expressed or spliced in the transition to inflexible behavior. Splicing analysis suggests that repressed activity in the DMS during the transition to inflexible behavior reflects both reduced expression and increased degradation of plasticity-related mRNA transcripts. Finally, given the ability of CORT to influence histone acetylation, we defined CORT-mediated H3K9ac enrichment profiles associated with synaptic plasticity gene regulation stratified by sex and striatal subregion. This study is the first to define CORT-driven epigenetic regulation in the DMS and DLS during the transition from flexible to inflexible behavior in male and female mice.

## Introduction

Goal-directed behaviors are motivated by the expectation that actions will result in desirable outcomes. With repetition, such behaviors can take on inflexible, routinized qualities that are insensitive to outcome value [1–3]. Behavioral inflexibility allows organisms to conserve cognitive resources when a given behavior has been reliably reinforced in the past, but this can occur at the expense of adapting to unexpected or negative events. Accordingly, the loss of behavioral flexibility is thought to contribute to psychopathologies, including obsessive-compulsive disorders, addiction, binging disorders, and neurodegenerative diseases [4–9].

The transition from flexible to inflexible behavior naturally occurs with extended experience with a given task and is well characterized and regulated by specific corticostriatal afferents [2,3]. Most critically, the dorsomedial striatum (DMS) is necessary for producing flexible behavior [1–3,10], while the dorsolateral striatum (DLS) is a primary driver of inflexible behavior [1–3,11]. Still, molecular mediators are only beginning to be revealed. The present study aimed to define gene regulation that underlies the transition from behavioral flexibility to inflexibility.

Stress influences decision-making behavior [12,13]. While acute stress can heighten vigilance and focus, promoting adaptive action [14,15], chronic stress often impairs cognition, resulting in suboptimal decision-making [14–16] and inflexible response bias [17–19]. The glucocorticoid receptor (GR) is a primary stress hormone receptor in rodents [20–22], activated by the agonist corticosterone (CORT) in rodents and cortisol in humans [23]. Chronic elevated CORT is sufficient to drive the loss of flexible behaviors in male rodents [17,24–26]. In this study we expand these findings to female mice, and confirm that GR activation is necessary for CORT acceleration of behavioral inflexibility.

Stress-related changes in decision-making strategies are accompanied by dendritic atrophy on neurons in the DMS and dendritic outgrowth on neurons in the DLS [17]. The structural and morphological adaptations of neurons undergoing synaptic plasticity allow for the refinement of circuit outputs by increasing or decreasing the strength of neuronal connections within or between brain regions [27–29]. We discovered that CORT-accelerated loss of behavioral flexibility is accompanied by synaptic plasticity-related gene expression or alternative splicing in the DMS and DLS. This study is a comprehensive examination of sex- and striatal-subregion-specific gene expression, alternative splicing, and histone acetylation during the loss of behavioral flexibility following extended training or CORT exposure.

## Materials and Methods

### Animals

8-week-old male and female C57BL/6J mice were purchased from The Jackson Laboratory (https://www.jax.org/strain/000664) and were housed under a 12-hour light-dark cycle (lights on 7:00 AM) at 23 °C. Mice were allowed a week to habituate, then calorie-restricted to maintain 90 ± 5% of their individual baseline body weight for the remainder of the experiments. Mice were housed in same-sex cages with two to four age- and sex-matched cagemates. All experiments were performed in accordance with the University of Pennsylvania Institutional Animal Care and Use Committee and were conducted in accordance with the National Institute of Health Guide for the Care and Use of Laboratory Animals.

### Statistical analysis for behavioral studies

Data are expressed as mean ± standard error of the mean (SEM). Statistical analyses were performed using GraphPad Prism software (version 10.2.2, La Jolla, CA). For all behavioral analyses, we employed an ANOVA followed by a Tukey or Šidák post-hoc test with an alpha of 0.05 to correct for multiple comparisons. Sex as a biological variable was considered in the statistical analyses of all behavioral results. Two-way ANOVA was used to compare two variables (i.e., sex and training), and post-hoc comparisons followed interaction effects. Three-way ANOVA was used to compare three variables (i.e., sex, CORT, and training), with post-hoc comparisons following interaction effects. Full F statistics are reported for ANOVAs, while degrees of freedom (DF) are reported for post-hoc comparisons. The “training day” variable was included as a repeated-measure across all operant conditioning sessions to analyze food port entry rates and lever pressing rates for individual mice undergoing multiple days of operant sessions. The “training duration” variable was included as a discrete variable (limited or extended) when analyzing testing session data, as mice could only receive one pre-training before testing. The Grubbs test was used to identify outliers. Outlier mice were removed from all analyses without replacement. A table of all relevant statistical analyses for each Figure and Supplemental Figure has been provided, with all experimental variables included.

Detailed materials and methods is described in the supplemental materials.

## Results

### Chronic CORT administration accelerates the loss of behavioral flexibility

Operant conditioning occurs when mice engage in a behavior (e.g. lever pressing) to earn a reinforcement (e.g. sucrose). A limited number of operant training days results in flexible operant behavior, which is sensitive to changes in reinforcement value or availability. Extended training causes motor sequences to become routinized, resulting in inflexible behavior, which is insensitive to changes in reinforcement value [30,31]. Stress exposure promotes inflexible behavior through activation of the glucocorticoid receptor (GR), a primary factor controlling the stress response [32], and plasma CORT increases with chronic stress [33,34]. To examine the molecular underpinnings of CORT-accelerated behavioral inflexibility, we trained both male and female mice in a standard protocol for operant conditioning under limited and extended training conditions [35–37], with or without CORT treatment **(Figures 1A-B, S1A-C)**. We assigned mice to either vehicle (1% ethanol) or CORT (50µg/mL) drinking water groups. The CORT dose was based on previous literature [24,25,38] and a dose-response study that confirmed an average CORT dose of 8mg/kg/day in male and female mice (**Figure 1B**; n = 12-13 mice/group, two-way ANOVA, main effect of Sex, F_(1, 23)_ = 1.842, *p = 0.1878*).

**Figure 1:**
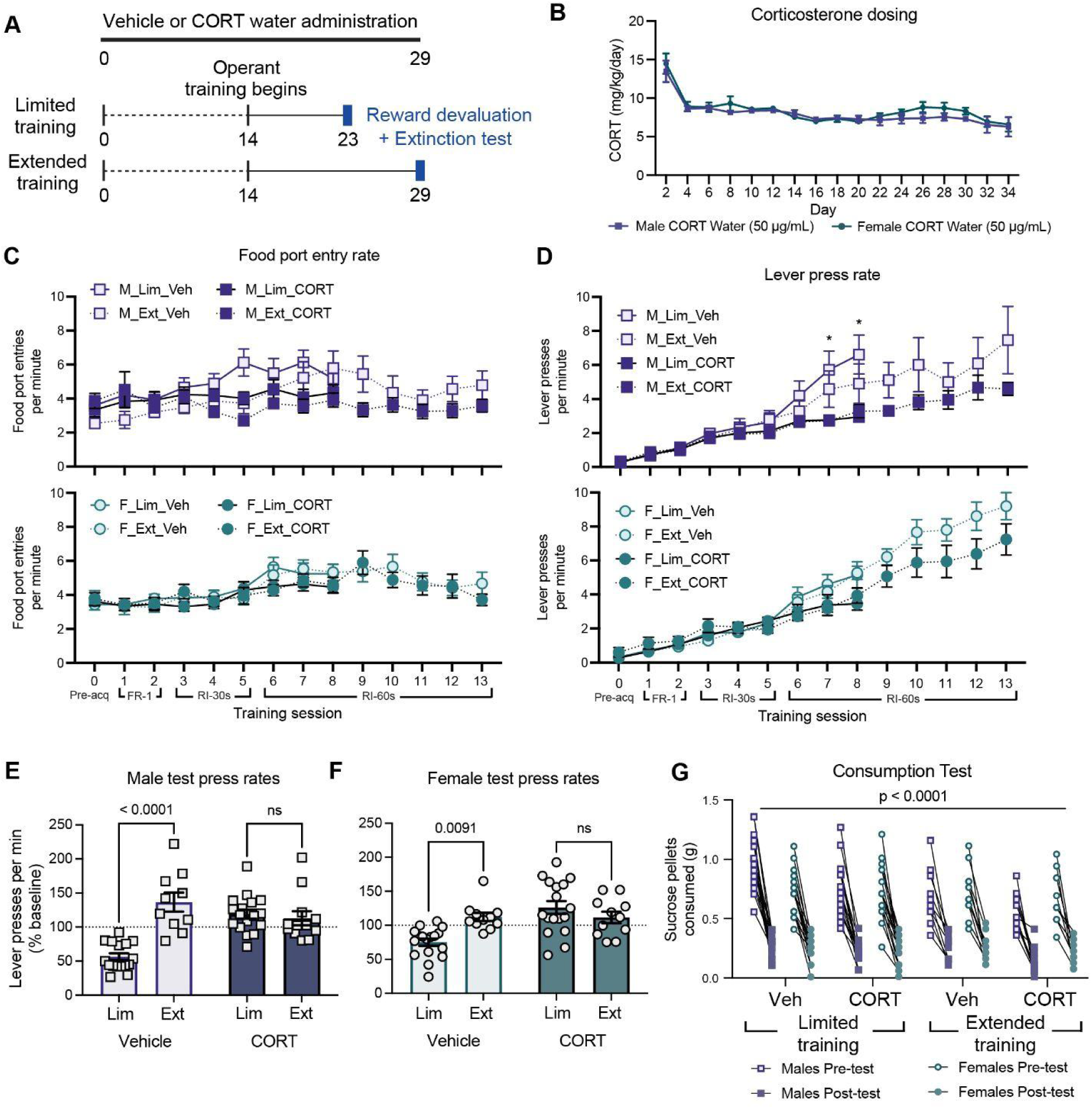
Chronic CORT administration accelerates the loss of behavioral flexibility. **A** Behavioral timeline where CORT or vehicle water administration began 14 days before training and lasted throughout the duration of training and testing. All mice underwent two days of FR-1 training, three days of RI-30s training, then either three days of RI-60s training for limited training groups, or eight days of RI-60s training for extended training groups. Mice progressed to a reward devaluation and testing the day after their final RI-60s training session. **B** There was no sex difference in CORT dosing when adjusting for body weight and water intake per cage [n = 12-13 cages/sex, two-way ANOVA, main effect of Time, F_(16, 296)_ = 18.36, *p < 0.0001*; main effect of Sex, F_(1, 23)_ = 1.842, *p = 0.1878*]. **C, D** The average food port entry rate per minute or lever pressing rate per minute ± SEM by sex (male or female), training duration (limited or extended), and treatment (vehicle or CORT) is shown across each day of the operant conditioning task. CORT had no effect on food port entries in limited [n = 16-17 mice/group, three-way ANOVA, main effect of Training Day, F_(8, 495)_ = 10.96, *p < 0.0001*; main effect of Sex, F_(1, 62)_ = 1.147, *p = 0.2884*; main effect of CORT, F_(1, 62)_ = 3.515, *p = 0.0655*] or extended groups [n = 10-12 mice/group, three-way ANOVA, main effect of Training Day, F_(13, 516)_ = 6.474, *p < 0.0001*; main effect of Sex, F_(1, 40)_ = 2.652, *p = 0.1113*; main effect of CORT, F_(1, 40)_ = 1.495, *p = 0.2286*]. CORT decreased the lever pressing rates in limited [n = 16-17 mice/group, three-way ANOVA, main effect of Training Day, F_(8, 423)_ = 76.77, *p < 0.0001*; main effect of Sex, F_(1, 53)_ = 0.2827, *p = 0.5971*; main effect of CORT, F_(1, 53)_ = 6.999, *p = 0.0107*] and extended groups [n = 10-12 mice/group, three-way ANOVA, main effect of Training Day, F_(13, 516)_ = 83.37, *p < 0.0001*; main effect of Sex, F_(1, 40)_ = 3.321, *p = 0.0759*; main effect of CORT, F_(1, 40)_ = 4.346, *p = 0.0435*]. All mice exhibited an increase in lever pressing as the random interval for training increased. **E, F** In an unrewarded lever pressing test following sensory-specific satiety devaluation, vehicle extended training and both CORT training groups do not attenuate lever pressing and display inflexible lever pressing [n = 10-17 mice/group, three-way ANOVA followed by Tukey post hoc tests, main effect of Training Duration, F_(1, 52)_ = 0.7872, *p < 0.0001*; main effect of Sex, F_(1, 52)_ = 12.85, *p = 0.987*; main effect of CORT, F_(1, 52)_ = 38.91, *p = 0.0002*; VehMalesLimited v VehMalesExtended, DF = 103, *p < 0.0001*; VehMalesLimited v CORTMalesLimited, DF = 103, *p < 0.0001*; VehMalesLimited v CORTMalesExtended, DF = 103, *p < 0.0001*; VehFemalesLimited v VehFemalesExtended, DF = 103, *p = 0.0091*; VehFemalesLimited v CORTFemalesLimited, DF = 103, *p < 0.0001*; VehFemalesLimited v CORTFemalesExtended, DF = 103, *p = 0.0478*]. Lever presses per minute from the test are reported as the percent of an individual’s baseline from the average of the individual’s last two completed training sessions. **G** All mice reduced free consumption of the sucrose reward following testing compared to their pre-test consumption during sensory-specific satiety devaluation [n = 10-17 mice/group, three-way ANOVA, main effect of Pre v Post Consumption Test, F_(1, 107)_ = 594.4, *p < 0.0001*]. See **Table 1** for detailed statistics for Figure 1. See **Figure S1** for detailed plasma CORT levels and **Table S1** for subsequent statistics.

**Table 1:** Detailed statistics for. **Figure 1**. Either two-way or three-way analysis of the variance (ANOVA) tests were run with either a Šidák or Tukey post-hoc to correct for multiple comparisons. Corresponding figures, metrics, ANOVA comparisons, F statistics, and p values are shown.

| Figure | Metric | ANOVA comparison | F(DFn, DFd) | p value |
| --- | --- | --- | --- | --- |
| Fig. 1B | Corticosterone dosing | Time | F(16, 296) = 18.36 | < 0.0001 |
|  |  | Sex | F(1, 23) = 1.842 | 0.1878 |
|  |  | Time x Sex | F(16, 296) = 0.5286 | 0.9313 |
| Fig. 1C | Food port entry rate - limited training | Training Day | F (8, 495) = 10.96 | < 0.0001 |
|  |  | Sex | F (1, 62) = 1.147 | 0.2884 |
|  |  | CORT | F (1, 62) = 3.515 | 0.0655 |
|  |  | Training Day x Sex | F (8, 495) = 1.664 | 0.1047 |
|  |  | Training Day x CORT | F (8, 495) = 1.363 | 0.2102 |
|  |  | Sex x CORT | F (1, 62) = 0.3044 | 0.5831 |
|  |  | Training Day x Sex x CORT | F (8, 495) = 1.082 | 0.3744 |
| Fig. 1C | Food port entry rate - limited training | Males: Vehicle v CORT D0 (Šídák post-hoc) | DF = 476 | > 0.9999 |
|  |  | Males: Vehicle v CORT D1 (Šídák post-hoc) | DF = 476 | > 0.9999 |
|  |  | Males: Vehicle v CORT D2 (Šídák post-hoc) | DF = 476 | > 0.9999 |
|  |  | Males: Vehicle v CORT D3 (Šídák post-hoc) | DF = 476 | > 0.9999 |
|  |  | Males: Vehicle v CORT D4 (Šídák post-hoc) | DF = 476 | > 0.9999 |
|  |  | Males: Vehicle v CORT D5 (Šídák post-hoc) | DF = 476 | 0.8445 |
|  |  | Males: Vehicle v CORT D6 (Šídák post-hoc) | DF = 476 | > 0.9999 |
|  |  | Males: Vehicle v CORT D7 (Šídák post-hoc) | DF = 476 | 0.9972 |
|  |  | Males: Vehicle v CORT D8 (Šídák post-hoc) | DF = 476 | < 0.0001 |
| Fig. 1C | Food port entry rate - limited training | Females: Vehicle v CORT D0 (Šídák post-hoc) | DF = 476 | > 0.9999 |
|  |  | Females: Vehicle v CORT D1 (Šídák post-hoc) | DF = 476 | > 0.9999 |
|  |  | Females: Vehicle v CORT D2 (Šídák post-hoc) | DF = 476 | > 0.9999 |
|  |  | Females: Vehicle v CORT D3 (Šídák post-hoc) | DF = 476 | > 0.9999 |
|  |  | Females: Vehicle v CORT D4 (Šídák post-hoc) | DF = 476 | > 0.9999 |
|  |  | Females: Vehicle v CORT D5 (Šídák post-hoc) | DF = 476 | > 0.9999 |
|  |  | Females: Vehicle v CORT D6 (Šídák post-hoc) | DF = 476 | > 0.9999 |
|  |  | Females: Vehicle v CORT D7 (Šídák post-hoc) | DF = 476 | > 0.9999 |
|  |  | Females: Vehicle v CORT D8 (Šídák post-hoc) | DF = 476 | > 0.9999 |
| **Fig. 1C** | Food port entry rate - extended training | Training Day | F (13, 516) = 6.474 | < 0.0001 |
|  |  | Sex | F (1, 40) = 2.652 | 0.1113 |
|  |  | CORT | F (1, 40) = 1.495 | 0.2286 |
|  |  | Training Day x Sex | F (13, 516) = 1.180 | 0.2906 |
|  |  | Training Day x CORT | F (13, 516) = 2.513 | 0.0024 |
|  |  | Sex x CORT | F (1, 40) = 0.2365 | 0.6294 |
|  |  | Training Day x Sex x CORT | F (13, 516) = 1.673 | 0.063 |
| **Fig. 1C** | Food port entry rate - extended training | Males: Vehicle v CORT D0 (Šídák post-hoc) | DF = 556 | > 0.9999 |
|  |  | Males: Vehicle v CORT D1 (Šídák post-hoc) | DF = 556 | > 0.9999 |
|  |  | Males: Vehicle v CORT D2 (Šídák post-hoc) | DF = 556 | > 0.9999 |
|  |  | Males: Vehicle v CORT D3 (Šídák post-hoc) | DF = 556 | > 0.9999 |
|  |  | Males: Vehicle v CORT D4 (Šídák post-hoc) | DF = 556 | > 0.9999 |
|  |  | Males: Vehicle v CORT D5 (Šídák post-hoc) | DF = 556 | > 0.9999 |
|  |  | Males: Vehicle v CORT D6 (Šídák post-hoc) | DF = 556 | > 0.9999 |
|  |  | Males: Vehicle v CORT D7 (Šídák post-hoc) | DF = 556 | > 0.9999 |
|  |  | Males: Vehicle v CORT D8 (Šídák post-hoc) | DF = 556 | > 0.9999 |
|  |  | Males: Vehicle v CORT D9 (Šídák post-hoc) | DF = 556 | 0.9999 |
|  |  | Males: Vehicle v CORT D10 (Šídák post-hoc) | DF = 556 | > 0.9999 |
|  |  | Males: Vehicle v CORT D11 (Šídák post-hoc) | DF = 556 | > 0.9999 |
|  |  | Males: Vehicle v CORT D12 (Šídák post-hoc) | DF = 556 | > 0.9999 |
|  |  | Males: Vehicle v CORT D13 (Šídák post-hoc) | DF = 556 | > 0.9999 |
| **Fig. 1C** | Food port entry rate - extended training | Females: Vehicle v CORT D0 (Šídák post-hoc) | DF = 556 | > 0.9999 |
|  |  | Females: Vehicle v CORT D1 (Šídák post-hoc) | DF = 556 | > 0.9999 |
|  |  | Females: Vehicle v CORT D2 (Šídák post-hoc) | DF = 556 | > 0.9999 |
|  |  | Females: Vehicle v CORT D3 (Šídák post-hoc) | DF = 556 | > 0.9999 |
|  |  | Females: Vehicle v CORT D4 (Šídák post-hoc) | DF = 556 | > 0.9999 |
|  |  | Females: Vehicle v CORT D5 (Šídák post-hoc) | DF = 556 | > 0.9999 |
|  |  | Females: Vehicle v CORT D6 (Šídák post-hoc) | DF = 556 | > 0.9999 |
|  |  | Females: Vehicle v CORT D7 (Šídák post-hoc) | DF = 556 | > 0.9999 |
|  |  | Females: Vehicle v CORT D8 (Šídák post-hoc) | DF = 556 | > 0.9999 |
|  |  | Females: Vehicle v CORT D9 (Šídák post-hoc) | DF = 556 | > 0.9999 |
|  |  | Females: Vehicle v CORT D10 (Šídák post-hoc) | DF = 556 | > 0.9999 |
|  |  | Females: Vehicle v CORT D11 (Šídák post-hoc) | DF = 556 | > 0.9999 |
|  |  | Females: Vehicle v CORT D12 (Šídák post-hoc) | DF = 556 | > 0.9999 |
|  |  | Females: Vehicle v CORT D13 (Šídák post-hoc) | DF = 556 | > 0.9999 |
| **Fig. 1D** | Lever pressing rate - limited training | Training Day | F (8, 423) = 76.77 | < 0.0001 |
|  |  | Sex | F (1, 53) = 0.2827 | 0.5971 |
|  |  | CORT | F (1, 53) = 6.999 | 0.0107 |
|  |  | Training Day x Sex | F (8, 423) = 0.2266 | 0.986 |
|  |  | Training Day x CORT | F (8, 423) = 8.977 | < 0.0001 |
|  |  | Sex x CORT | F (1, 53) = 1.362 | 0.2484 |
|  |  | Training Day x Sex x CORT | F (8, 423) = 1.217 | 0.2872 |
| **Fig. 1D** | Lever pressing rate - limited training | Males: Vehicle v CORT D0 (Šídák post-hoc) | DF = 476 | > 0.9999 |
|  |  | Males: Vehicle v CORT D1 (Šídák post-hoc) | DF = 476 | > 0.9999 |
|  |  | Males: Vehicle v CORT D2 (Šídák post-hoc) | DF = 476 | > 0.9999 |
|  |  | Males: Vehicle v CORT D3 (Šídák post-hoc) | DF = 476 | > 0.9999 |
|  |  | Males: Vehicle v CORT D4 (Šídák post-hoc) | DF = 476 | > 0.9999 |
|  |  | Males: Vehicle v CORT D5 (Šídák post-hoc) | DF = 476 | > 0.9999 |
|  |  | Males: Vehicle v CORT D6 (Šídák post-hoc) | DF = 476 | > 0.9999 |
|  |  | Males: Vehicle v CORT D7 (Šídák post-hoc) | DF = 476 | 0.0042 |
|  |  | Males: Vehicle v CORT D8 (Šídák post-hoc) | DF = 476 | < 0.0001 |
| **Fig. 1D** | Lever pressing rate - limited training | Females: Vehicle v CORT D0 (Šídák post-hoc) | DF = 476 | > 0.9999 |
|  |  | Females: Vehicle v CORT D1 (Šídák post-hoc) | DF = 476 | > 0.9999 |
|  |  | Females: Vehicle v CORT D2 (Šídák post-hoc) | DF = 476 | > 0.9999 |
|  |  | Females: Vehicle v CORT D3 (Šídák post-hoc) | DF = 476 | > 0.9999 |
|  |  | Females: Vehicle v CORT D4 (Šídák post-hoc) | DF = 476 | > 0.9999 |
|  |  | Females: Vehicle v CORT D5 (Šídák post-hoc) | DF = 476 | > 0.9999 |
|  |  | Females: Vehicle v CORT D6 (Šídák post-hoc) | DF = 476 | > 0.9999 |
|  |  | Females: Vehicle v CORT D7 (Šídák post-hoc) | DF = 476 | > 0.9999 |
|  |  | Females: Vehicle v CORT D8 (Šídák post-hoc) | DF = 476 | 0.6345 |
| **Fig. 1D** | Lever pressing rate - extended training | Training Day | F (13, 516) = 83.37 | < 0.0001 |
|  |  | Sex | F (1, 40) = 3.321 | 0.0759 |
|  |  | CORT | F (1, 40) = 4.346 | 0.0435 |
|  |  | Training Day x Sex | F (13, 516) = 4.321 | < 0.0001 |
|  |  | Training Day x CORT | F (13, 516) = 4.045 | < 0.0001 |
|  |  | Sex x CORT | F (1, 40) = 0.1337 | 0.7165 |
|  |  | Training Day x Sex x CORT | F (13, 516) = 0.3406 | 0.9853 |
| **Fig. 1D** | Lever pressing rate - extended training | Males: Vehicle v CORT D0 (Šídák post-hoc) | DF = 556 | > 0.9999 |
|  |  | Males: Vehicle v CORT D1 (Šídák post-hoc) | DF = 556 | > 0.9999 |
|  |  | Males: Vehicle v CORT D2 (Šídák post-hoc) | DF = 556 | > 0.9999 |
|  |  | Males: Vehicle v CORT D3 (Šídák post-hoc) | DF = 556 | > 0.9999 |
|  |  | Males: Vehicle v CORT D4 (Šídák post-hoc) | DF = 556 | > 0.9999 |
|  |  | Males: Vehicle v CORT D5 (Šídák post-hoc) | DF = 556 | > 0.9999 |
|  |  | Males: Vehicle v CORT D6 (Šídák post-hoc) | DF = 556 | > 0.9999 |
|  |  | Males: Vehicle v CORT D7 (Šídák post-hoc) | DF = 556 | > 0.9999 |
|  |  | Males: Vehicle v CORT D8 (Šídák post-hoc) | DF = 556 | > 0.9999 |
|  |  | Males: Vehicle v CORT D9 (Šídák post-hoc) | DF = 556 | > 0.9999 |
|  |  | Males: Vehicle v CORT D10 (Šídák post-hoc) | DF = 556 | > 0.9999 |
|  |  | Males: Vehicle v CORT D11 (Šídák post-hoc) | DF = 556 | > 0.9999 |
|  |  | Males: Vehicle v CORT D12 (Šídák post-hoc) | DF = 556 | > 0.9999 |
|  |  | Males: Vehicle v CORT D13 (Šídák post-hoc) | DF = 556 | 0.8447 |
| **Fig. 1D** | Lever pressing rate - extended training | Females: Vehicle v CORT D0 (Šídák post-hoc) | DF = 556 | > 0.9999 |
|  |  | Females: Vehicle v CORT D1 (Šídák post-hoc) | DF = 556 | > 0.9999 |
|  |  | Females: Vehicle v CORT D2 (Šídák post-hoc) | DF = 556 | > 0.9999 |
|  |  | Females: Vehicle v CORT D3 (Šídák post-hoc) | DF = 556 | > 0.9999 |
|  |  | Females: Vehicle v CORT D4 (Šídák post-hoc) | DF = 556 | > 0.9999 |
|  |  | Females: Vehicle v CORT D5 (Šídák post-hoc) | DF = 556 | > 0.9999 |
|  |  | Females: Vehicle v CORT D6 (Šídák post-hoc) | DF = 556 | > 0.9999 |
|  |  | Females: Vehicle v CORT D7 (Šídák post-hoc) | DF = 556 | > 0.9999 |
|  |  | Females: Vehicle v CORT D8 (Šídák post-hoc) | DF = 556 | > 0.9999 |
|  |  | Females: Vehicle v CORT D9 (Šídák post-hoc) | DF = 556 | > 0.9999 |
|  |  | Females: Vehicle v CORT D10 (Šídák post-hoc) | DF = 556 | > 0.9999 |
|  |  | Females: Vehicle v CORT D11 (Šídák post-hoc) | DF = 556 | > 0.9999 |
|  |  | Females: Vehicle v CORT D12 (Šídák post-hoc) | DF = 556 | > 0.9999 |
|  |  | Females: Vehicle v CORT D13 (Šídák post-hoc) | DF = 556 | > 0.9999 |
| **Fig. 1E, F** | Male and Female test press rates | Training Duration | F (1, 52) = 0.7872 | < 0.0001 |
|  |  | Sex | F (1, 52) = 12.85 | 0.987 |
|  |  | CORT | F (1, 52) = 38.91 | 0.0002 |
|  |  | Training Duration x Sex | F (1, 52) = 0.005158 | 0.0292 |
|  |  | Training Duration x CORT | F (1, 52) = 0.4073 | < 0.0001 |
|  |  | Sex x CORT | F (1, 52) = 6.164 | 0.6316 |
|  |  | Training Duration x Sex x CORT | F (1, 52) = 9.935 | 0.1414 |
| **Fig. 1E, F** | Male and Female test press rates | Veh_Male_Limited training v Veh_Male_Extended training (Tukey post-hoc) | DF = 103 | < 0.0001 |
|  |  | Veh_Male_Limited training v CORT_Male_Limited training (Tukey post-hoc) | DF = 103 | < 0.0001 |
|  |  | Veh_Male_Limited training v CORT_Male_Extended training (Tukey post-hoc) | DF = 103 | < 0.0001 |
|  |  | CORT_Male_Limited training v Veh_Male_Extended training (Tukey post-hoc) | DF = 103 | 0.8207 |
|  |  | CORT_Male_Limited training v CORT_Male_Extended training (Tukey post-hoc) | DF = 103 | 0.9995 |
|  |  | Veh_Female_Limited training v Veh_Female_Extended training (Tukey post-hoc) | DF = 103 | 0.0091 |
|  |  | Veh_Female_Limited training v CORT_Female_Limited training (Tukey post-hoc) | DF = 103 | < 0.0001 |
|  |  | Veh_Female_Limited training v CORT_Female_Extended training (Tukey post-hoc) | DF = 103 | 0.0478 |
|  |  | CORT_Female_Limited training v<br>Veh_Female_Extended training<br>(Tukey post-hoc) | DF = 103 | 0.9333 |
|  |  | CORT_Female_Limited training v<br>CORT_Female_Extended training<br>(Tukey post-hoc) | DF = 103 | 0.9141 |
|  |  | Veh_Male_Limited training v<br>Veh_Female_Limited training (Tukey<br>post-hoc) | DF = 103 | < 0.0001 |
|  |  | CORT_Male_Limited training v<br>CORT_Female_Limited training<br>(Tukey post-hoc) | DF = 103 | 0.997 |
|  |  | Veh_Male_Extended training v<br>Veh_Female_Extended training<br>(Tukey post-hoc) | DF = 103 | 0.5883 |
|  |  | CORT_Male_Extended training v<br>CORT_Female_Extended training<br>(Tukey post-hoc) | DF = 103 | > 0.9999 |
| **Fig. 1G** | Consumption test | Pre v Post | F(1, 107) = 594.4 | < 0.0001 |
|  |  | Sex | F(1, 107) = 0.6156 | 0.4344 |
|  |  | CORT | F(1, 107) = 9.567 | 0.0025 |
|  |  | Pre v Post x Sex | F(1, 107) = 2.705 | 0.103 |
|  |  | Pre v Post x CORT | F(1, 107) = 1.677 | 0.1981 |
|  |  | Sex x CORT | F(1, 107) = 6.795 | 0.2487 |
|  |  | Pre v Post x Sex x CORT | F(1, 107) = 2.401 | 0.1242 |

We first tested the hypothesis that CORT extinguishes behavioral flexibility in both male [24–26] and female mice. CORT or vehicle was administered for two weeks prior to and throughout training and testing (**Figure 1A**). To avoid modeling natural variation in pressing rates, mice were balanced to limited or extended groups based on days two and three of random interval (RI-)60s lever pressing rates. We confirmed that groups did not differ during FR-1, RI-30s, and through RI-60s day 3 (**Figures S2A-C**). Food port entry was reduced by CORT, with no main effect of sex (**Figure 1C**; n = 10-17 mice/group, three-way ANOVA, main effect of Training Day, F_(13, 1051)_ = 12.29, *p < 0.0001*; main effect of Sex, F_(1, 107)_ = 0.1941, *p = 0.6604*; main effect of CORT, F_(1, 107)_ = 7.502, *p = 0.0072*). Lever pressing increased across training days, as expected. CORT decreased lever pressing rates during training in both male and female mice, compared to vehicle (**Figure 1D**; n = 10-17 mice/group, three-way ANOVA, main effect of Training Day, F_(13, 1051)_ = 132.5, *p < 0.0001*; main effect of CORT, F_(1, 107)_ = 22.97, *p < 0.0001*; interaction effect of Training x CORT, F_(13, 1051)_ = 9.861, *p < 0.0001*), as previously reported with CORT-treated male mice [25]. There was no sex difference in lever pressing rates among vehicle or CORT groups (**Figure 1D**; n = 10-17 mice/group, three-way ANOVA, main effect of Sex, F_(1, 107)_ = 0.855, *p = 0.3572*; interaction effect of Sex x CORT, F_(1, 107)_ = 2.17, *p = 0.1437*).

Non-devalued limited and extended training male mice do not differ in lever pressing rates during an unrewarded probe test [35,37,39], even with chronic stress or CORT treatment [24–26]. Extended training male mice press more during an unrewarded probe test, indicating a transition to inflexible behavior [24–26,35,37,39]. We therefore hypothesized that limited operant training with CORT in both male and female mice would cause insensitivity to reward devaluation and behavioral inflexibility. We confirmed this with a standard paradigm previously applied only to male rodents. Following limited or extended training, mice underwent sucrose reward devaluation through satiety, followed by an unrewarded lever pressing test. As expected, mice that underwent limited training reduced lever pressing relative to their training baseline, demonstrating sensitivity to reduced reinforcer value. Alternatively, mice that underwent extended training did not integrate reinforcer value into their response strategies, with response rates equivalent to their training baseline (**Figure 1E**; n = 10-17 mice/group, three-way ANOVA followed by Tukey post-hoc tests, VehMaleLimitedTraining v VehMaleExtendedTraining, DF = 103, *p < 0.0001*; **Figure 1F**; n = 10-17 mice/group, three-way ANOVA followed by Tukey post-hoc tests, VehFemaleLimitedTraining v VehFemaleExtendedTraining, DF = 103, *p = 0.0131*; **Figures S3A–H**). Furthermore, CORT-treated mice with limited training did not exhibit sensitivity to reinforcer value, responding as during training and to the same extent as the extended training groups (**Figure 1E**; n = 10-17 mice/group, three-way ANOVA followed by Tukey post-hoc tests, VehMaleLimitedTraining v CORTMaleLimitedTraining, DF = 103, *p < 0.0001*; CORTMaleLimitedTraining v VehMaleExtendedTraining, DF = 103, *p = 0.7106*; **Figure 1F**; n = 10-17 mice/group, three-way ANOVA followed by Tukey post-hoc tests, VehFemaleLimitedTraining v CORTFemaleLimitedTraining, DF = 103, *p < 0.0001*; CORTFemaleLimitedTraining v VehFemaleExtendedTraining, DF = 103, *p = 0.9946*).Insensitivity to reinforcer value was not due to a failure to devalue the sucrose, as all mice consumed less sucrose in the post-probe consumption test (**Figure 1G**; n = 10-17 mice/group, three-way ANOVA, main effect of Pre v Post Consumption Test, F_(1, 107)_ = 594.4, *p < 0.0001*). These differences in testing press rates were also not attributable to differences in baseline training press rates (**Figures S2B–C**). Taken together, we showed that CORT is sufficient to extinguish behavioral flexibility in both male and female mice. Finally, we tested the hypothesis that GR activation is necessary for CORT-accelerated loss of behavioral flexibility. We found that co-administering CORT with the GR antagonist, mifepristone (MIF), during limited training rescued CORT-accelerated loss of behavioral flexibility in both male and female mice (**Figures S4A–H**).

### Dorsal striatum plasticity gene expression signatures during the CORT-mediated loss of behavioral flexibility

The dorsomedial striatum (DMS) is necessary for producing flexible behavior [1–3,10], while the dorsolateral striatum (DLS) is a primary driver of inflexible behavior [1–3,11]. The role of each brain region in each action strategy is supported by behaviorally relevant changes in synaptic morphology and physiology. We thus hypothesized that plasticity-related gene expression in the DMS and DLS reflect flexible or inflexible behavior, respectively. Having established that CORT drives the loss of behavioral flexibility with limited training in male and female mice (**Figures 1E–F**), we next tested the hypothesis that chronic CORT decreases DMS plasticity gene expression following limited training (**Figure 2A**). To test this, we reproduced the operant training paradigm and performed RNA-sequencing on DMS or DLS collected one hour after the final RI60s training session (**Figure 2A; Figure S5A**). Sequencing results were quality controlled through principal component analyses (**Figure 2A; Figures S5B–E**). RNA-seq and Gene Set Enrichment Analysis (GSEA) for CORT versus vehicle groups (sex-, brain region-, and training duration-matched) found that CORT decreased enrichment for synapse organization, axon development, and other plasticity-related gene sets in the DMS, but not in the DLS, following limited training compared to vehicle (**Figure 2B**).

**Figure 2:**
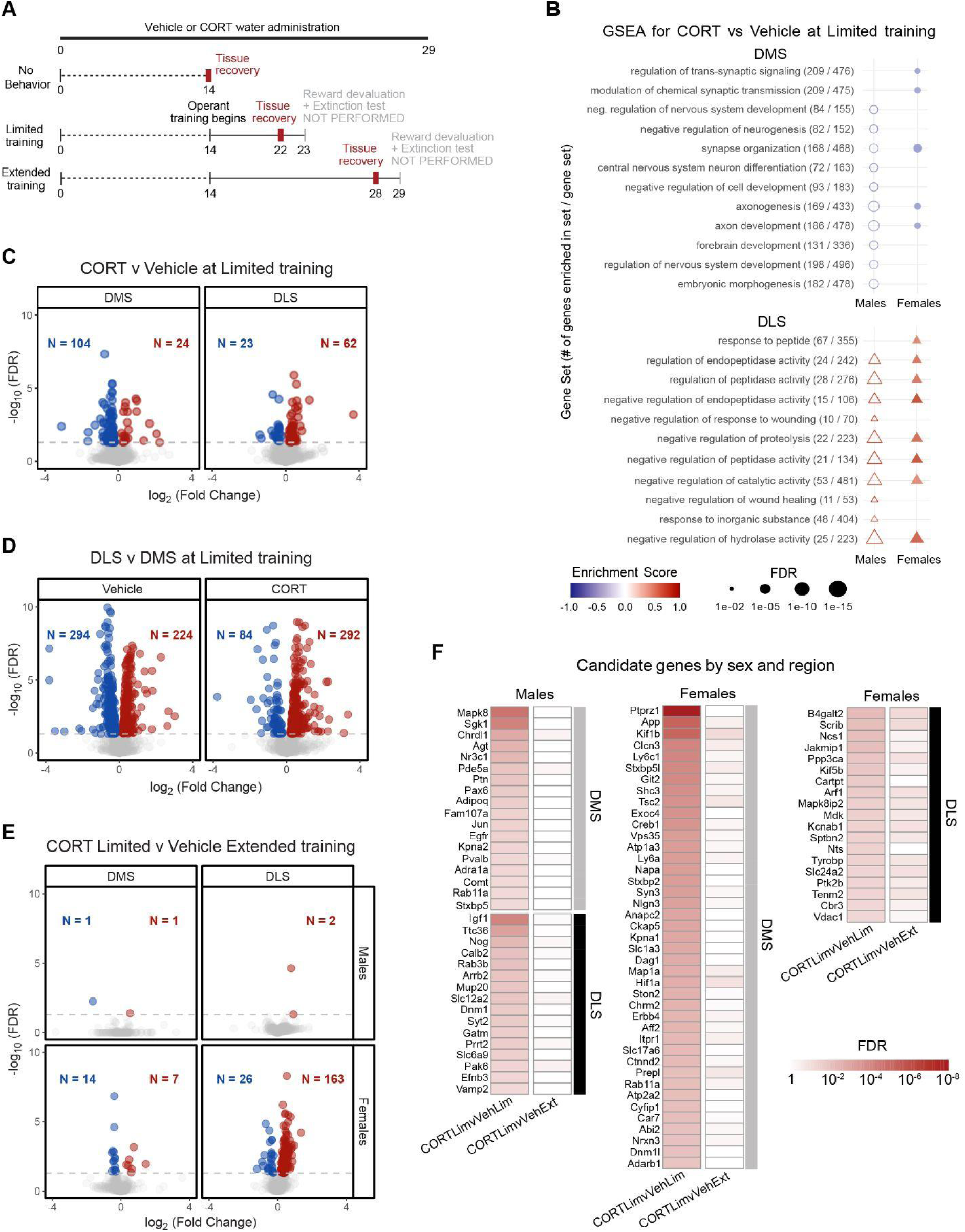
Dorsal striatum plasticity gene expression signatures during the CORT-mediated loss of behavioral flexibility. **A** Behavioral timeline for tissue recovery from No Behavior, Limited training, and Extended training time points. **B** A gene set enrichment analysis (GSEA) based on all differential gen e expression was performed for CORT versus vehicle groups after limited training. An independent GSEA was performed for each sex and brain region combination (n = 3/sex/region/treatment/training). The top ten GSEA terms by false discovery rate (FDR; Benjamini-Hochberg adjusted p-value) are shown for each combination of sex and brain region. The enrichment score is a scaling of the relative enrichment (red) or de-enrichment (blue) of a gene set term. Selection criteria included the top ten regulation terms per sex and region combination that have an FDR < 0.05. DLS terms were positively enriched for cellular activity regulation terms. DMS terms were negatively enriched for plasticity-relat ed terms. See **Figure S5** for tissue recovery locations and RNAseq principal component analyses. **C – E** Differentially expressed genes (DEGs) across 1157 genes from gene ontology terms: “synaptic signaling” and “cognition.” Significance cutoff (horizontal line at FDR = 0.05). **C** Differential gene expression from CORT versus vehicle groups produces a decrease in plasticity gene expression in the DMS and an increase in plasticity gene expression in the DLS. See **Figure S6** for detailed volcano plots. **D** Differential gene expression from DLS versus DMS groups produces similar amounts of up- and downregulated genes in vehicle groups, and produces less downregulation and more upregulation in CORT-treated groups. See **Figure S7** for detailed volcano plots. **E** Differential gene expression from CORT-limited versus vehicle-extended groups produces low differences in male DMS and DLS plasticity gene expression, while producing higher differences in female DMS and DLS plasticity gene expression. See **Figure S9** for detailed volcano plots. **F** Candidate genes by sex and region were determined by comparing the p.adjust values from the differential expression (DESEQ2) of the CORT limited training versus vehicle limited training groups (CORTLimvVehLim) against the false discovery rate (FDR) values from the CORT limited training versus vehicle extended training groups (CORTLimvVehExt). Genes that were changed in the CORTLimvVehLim comparison (FDR < 0.05) and were also not changed in the CORTLimvVehExt comparison (FDR > 0.05) are shown.

To deepen our understanding of decreased plasticity-related gene set enrichment in the DMS, we next examined gene expression at the individual gene level. Using an approach orthogonal to GSEA, we defined a list of 1157 plasticity-related genes included in the gene ontology (GO) terms, “synaptic signaling” and “cognition.” We selected these GO terms because they are significantly enriched in a meta-analysis of stress-regulated gene expression across behavioral paradigms [40]. In CORT versus vehicle groups, we found that chronic CORT downregulated expression of plasticity-related genes following limited training in the DMS (**Figure S6A**), in concordance with the GSEA outcomes **(Figure 2B)**. Next, we measured the expression of plasticity-related genes in the DLS, finding that chronic CORT increased plasticity gene expression in this region compared to vehicle (**Figure 2C; Figure S6B**). Specifically, in the DLS, CORT upregulated ∼3x more plasticity genes than it downregulated; while in the DMS, CORT downregulated ∼4x more plasticity genes than it upregulated **(Figure 2C)**. This ratio of CORT-influenced subregion-specific differential expression in DMS and DLS persisted into extended training (**Figures S6C–D**). When comparing DLS versus DMS plasticity gene expression, CORT downregulated plasticity DEGs and upregulated greater DEGs compared to vehicle following limited training (**Figure 2C; Figures S7A–B**), and these changes persisted through extended training (**Figures S7C–D**). Together, these analyses support our hypothesis that chronic CORT reduces DMS plasticity gene expression while increasing DLS plasticity gene expression following limited training compared to vehicle groups.

We found that the loss of behavioral flexibility following extended training was recapitulated by CORT treatment following limited training (**Figures 1E–F**). Additionally, when comparing vehicle extended to vehicle limited training groups, GSEA found reductions in plasticity gene set enrichment in the DMS (**Figure S8A**) with greater reduction in plasticity gene expression in the female DMS (**Figure S8B**). We thus tested the hypothesis that plasticity gene expression during vehicle extended training was recapitulated during CORT limited training in males. Indeed, CORT-limited versus vehicle-extended gene expression in males showed almost no differential gene expression (**Figure 2D; Figures S9A–B**). Alternatively, in females, gene expression differed between CORT-limited training and vehicle-extended training (**Figure 2D; Figures S9C–D**). These results suggest that plasticity-related gene expression during vehicle-extended training is recapitulated by limited training with CORT in male, but not female, mice.

We additionally sought to identify high-confidence candidate genes influenced by CORT that may be driving the loss of behavioral flexibility. To identify these candidate genes, we compiled a list of differentially expressed plasticity genes due to CORT by comparing CORT-limited training versus vehicle-limited training groups (CORTLimvVehLim). We then identified which of those plasticity genes were no longer differentially expressed when behavioral flexibility was lost in both groups (CORTLimvVehExt) (**Figure 2F**). This analysis provides the first list of candidate genes for direct and/or indirect CORT-influenced gene regulation in male and female DMS and DLS, and the uniqueness of the gene lists also suggests distinct gene expression responses to CORT and training by sex and region.

### Alternative splicing signatures during the loss of behavioral flexibility

Similar to gene expression, we hypothesized that CORT and training regulated differential alternative splicing in a region-specific manner during the loss of behavioral flexibility. Alternative splicing is an activity-dependent process that regulates transcript isoform abundance and produces functional protein differences such as differential stability, localization, binding properties, or enzymatic activity [41,42]. Importantly, neuronal activation regulates distinct genes by either differential alternative splicing or differential expression, requiring distinct bioinformatic analyses [43–45]. We first quantified alternative splicing events in each region, sex, and treatment condition. Next, we measured alternative splicing of plasticity-related genes as defined above. Critically, we found that plasticity-related genes were regulated by either differential expression or alternative splicing across sex and region (**Figure 3A**), such that distinct sets of plasticity genes are regulated at the level of total mRNA or the ratio of distinct isoforms.

**Figure 3:**
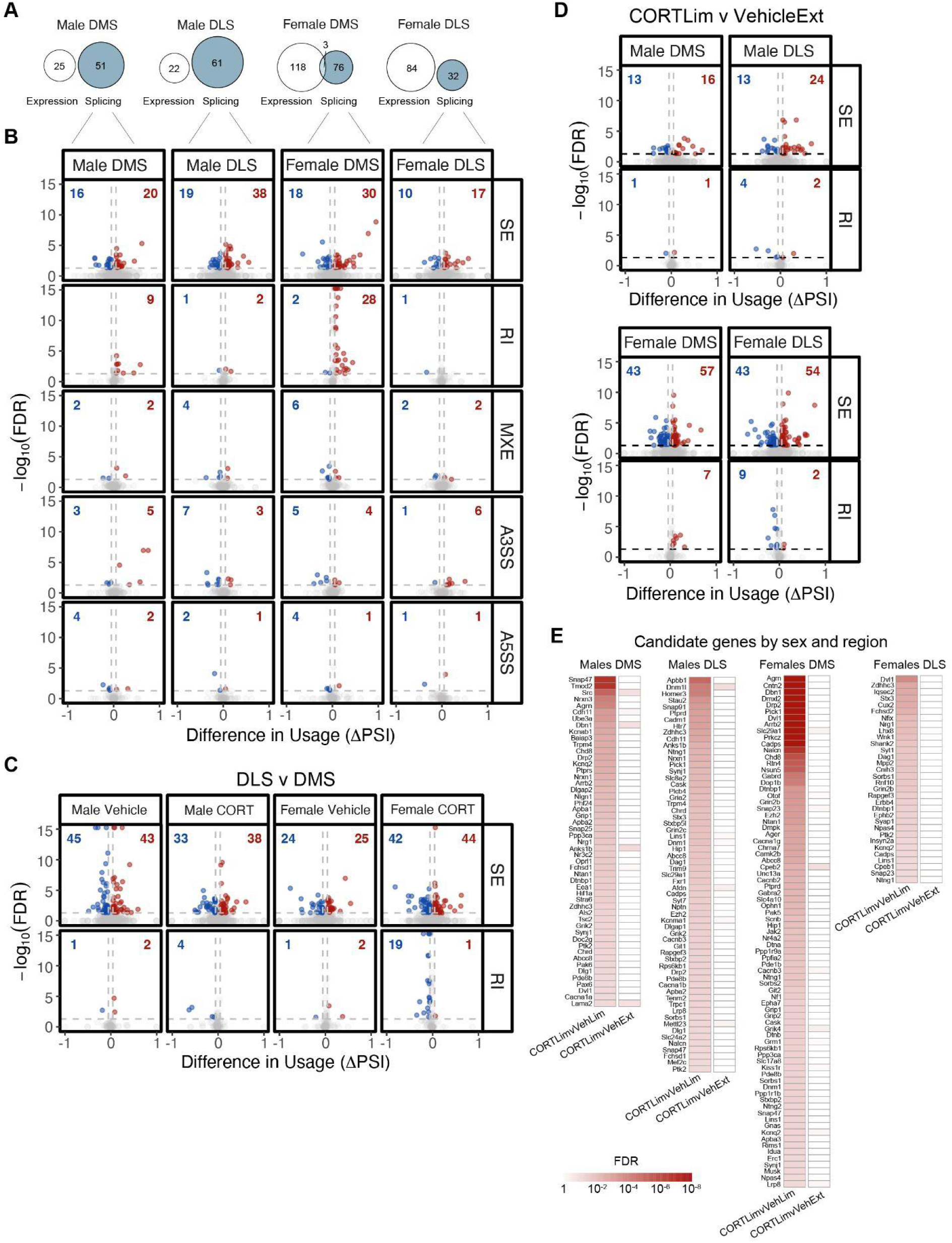
Alternative splicing signatures during the loss of behavioral flexibility. **A** 1157 genes were selected as plasticity-related genes from gene ontology terms: “synaptic signaling” and “cognition.” Of these genes, differential gene expression and differential splice events took place in distinct sets of plasticity genes across sex and region. **B** CORT administration increased or decreased the number of spliced exons (SEs) inclusions among plasticity genes following limited training. CORT administration increased the number of retained introns (RIs) in the DMS following limited training. There was no established trend for differences in mutually exclusive exons (MXE), alternative 3’ splice site (A3SS), or alternative 5’ splice site (A5SS). See **Figure S10** for relative proportions of SEs, RIs, MXEs, A5SSs, and A3SSs. **C** Differential inclusion of plasticity gene SEs was the largest difference between DLS and DMS across sex and treatment. The DLS had increased RIs in vehicle females, and the DLS had decreased RIs in male and female CORT groups. **D** There were considerable differences in SEs when comparing CORT-limited training to vehicle-extended groups. Additionally, the female DLS had fewer included RIs, and the female DMS had more included RIs when comparing CORT-limited training to vehicle-extended groups. The male DLS and DMS had relatively low changes in RIs by the same comparison. **E** Candidate genes by sex and region were determined by comparing the false discovery rate (FDR) values from the splice event difference in usage false discovery rate (FDR) of the CORT limited training versus vehicle limited training groups (CORTLimvVehLim) against the p.adjust values from the CORT limited training versus vehicle extended training groups (CORTLimvVehExt). Genes that were changed in the CORTLimvVehLim comparison (FDR < 0.05) and were also not changed in the CORTLimvVehExt comparison (FDR > 0.05) are shown.

Given that we observed region and sex-specific alternative splicing regulation of plasticity genes when comparing vehicle extended training to vehicle limited training (**Figure S8C**), we hypothesized that CORT regulates alternative splicing of plasticity-related genes. We quantified the differential inclusion of splice events in CORT versus vehicle groups following limited training across five separate events (skipped exon, SE, retained intron, RI, mutually exclusive exon, MXE, alternative 3’ splice site, A3SS, or alternative 5’ splice site, A5SS). Among plasticity-related genes, alternative SEs exhibited the greatest number of differential splicing events between CORT and vehicle groups following limited training, while the number of differential RI events was particularly distinct among DMS groups (**Figure 3B, Figure S10**). Comparing sex- and treatment-matched DLS versus DMS groups further suggested SEs were the most commonly disrupted splice event following limited training (**Figure 3C**). When comparing DLS versus DMS splice events, CORT treatment also produced a female-DLS-specific decrease (or female-DMS-specific increase) in RIs at the same limited training time point (**Figure 3C**). Our analyses suggest that SEs in the DMS and DLS are sensitive to CORT in both sexes, while RIs in the female DMS and DLS are more sensitive to CORT than in males.

As for differential expression, we hypothesized that alternative splicing of plasticity-related genes during extended training was recapitulated during CORT-limited training. When comparing CORTLimvVehExt training, we found both increased and decreased inclusion SEs in the DMS and DLS (**Figure 3D**). There were differential alternative RI events in the DMS and DLS in females, but not males (**Figure 3D**). Our analyses demonstrated that CORT, directly and/or indirectly, regulated total transcript levels or alternative splicing of distinct sets of plasticity-related genes following limited training.

We similarly sought to identify high-confidence candidate genes whose alternative splicing was influenced by CORT and may be driving the loss of behavioral flexibility. We used a similar approach to Figure 2F to identify which plasticity genes were differentially spliced due to CORT at limited training (CORTLimvVehLim) and no longer differentially expressed when comparing behaviorally inflexible groups (CORTLimvVehExt) (**Figure 3E**). This analysis provides the first list of candidate genes for direct and/or indirect CORT-influenced alternative splicing in male and female DMS and DLS. We find a greater number of commonly alternatively spliced genes across sex and region in response to CORT and training compared to the uniqueness of the differential expression candidates (**Figure 2E**).

### Sex- and region-specific contributions of H3K9ac to gene regulation during the CORT-mediated loss of behavioral flexibility

Given our confirmation of CORT-mediating the loss of behavioral flexibility in male and female mice [17,24,26], we next examined CORT-mediated chromatin regulation of plasticity-related gene expression. Acute GR activation recruits CREB-binding protein and p300 acetyltransferase to glucocorticoid response elements and increases global histone 3 lysine 9 acetylation (H3K9ac) enrichment in vitro [46,47], while chronic restraint stress reduces H3K9ac enrichment in mouse hippocampus [48]. Therefore, we hypothesized that GR activation via chronic CORT administration is sufficient to decrease global H3K9ac in mouse DMS and DLS. We measured global H3K9ac by ChIP-sequencing following limited training with CORT or vehicle in the DMS and DLS of the same mice from which we performed RNA-sequencing. We found H3K9ac enrichment in promoter, intragenic, and intergenic regions (**Figure 4A**). Given the relevance of promoter H3K9ac enrichment to gene expression, we restricted H3K9ac peak calling to promoter regions +/- 3kb from the transcription start site. Using aggregated H3K9ac promoter signal across all genes with called peaks, we compared CORTLimvVehLim training groups and found that chronic CORT decreased gene promoter H3K9ac enrichment globally compared to vehicle groups (**Figure 4B**), as expected.

**Figure 4:**
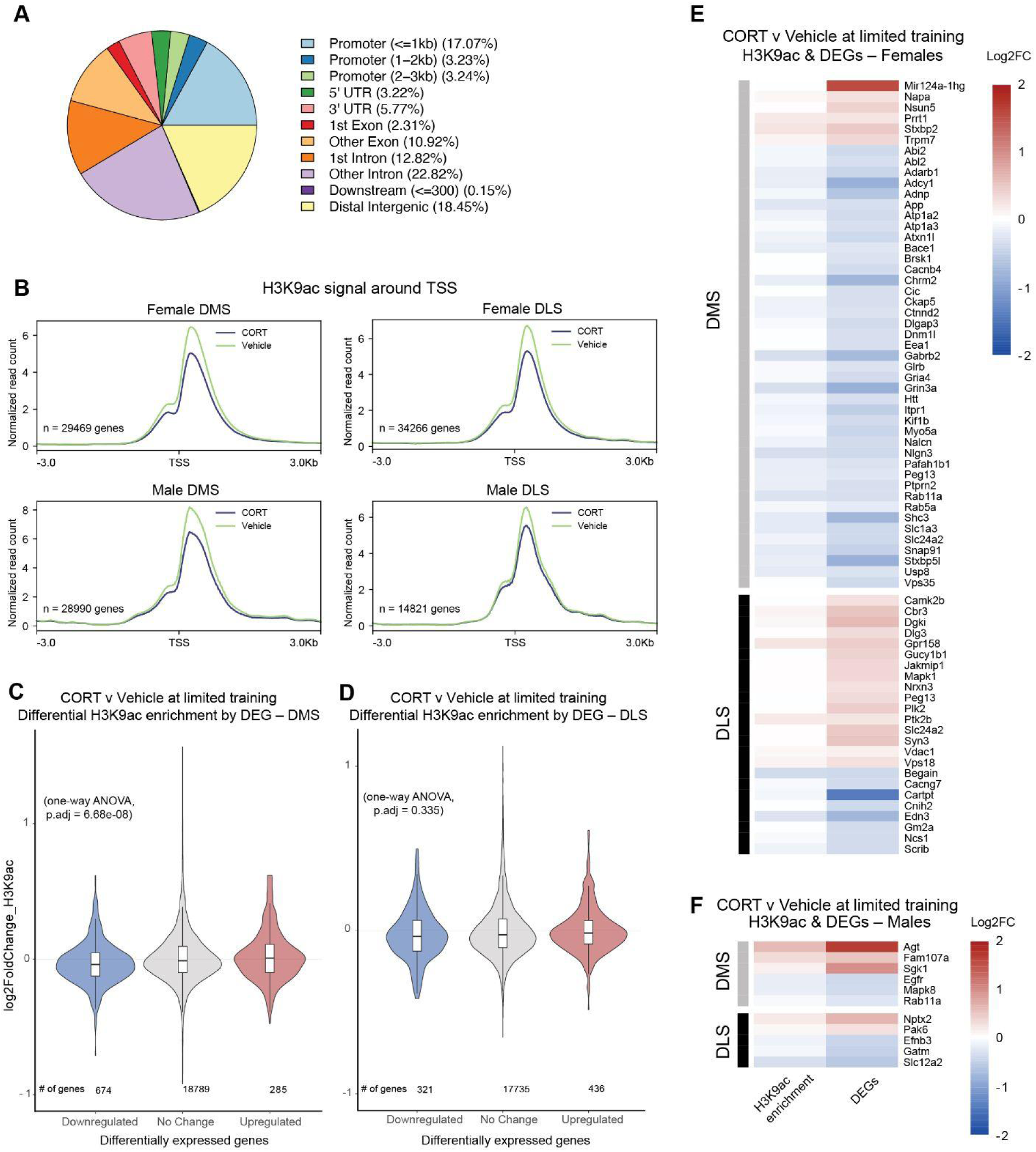
Sex- and region-specific contributions of H3K9ac to gene regulation during the CORT-mediated loss of behavioral flexibility. **A** H3K9ac enrichment distribution across all sample genomes (n = 3/sex/region/treatment/training). **B** CORT produces a global deenrichment of H3K9ac signal at all gene promoters centered around transcription start sites +/- 3kb for male and female DMS and DLS. **C, D** Upregulated and downregulated differentially expressed genes correlate with H3K9ac enrichment and deenrichment, respectively, in the DMS but not DLS. See **Table S5** for detailed statistics. See **Figure S11** for the extended training time point. **E, F** H3K9ac enrichment and deenrichment magnitude correlate with differentially expressed plasticity genes in female DMS and DLS, but not in male DMS and DLS.

Given our findings of region-specific differences in gene expression due to CORT administration (**Figures 2B–F**), we further hypothesized that CORT differentially enriched H3K9ac relevant to differential gene expression, in a brain region-specific manner following limited training. Differential H3K9ac enrichment analysis was performed between chronic CORT and vehicle groups disaggregated by sex and brain region to obtain log2 fold-change values for H3K9ac enrichment across all differentially expressed genes. We found a main effect of H3K9ac differential enrichment, such that H3K9ac enrichment from CORT versus vehicle groups positively correlated with differential gene expression in the DMS (**Figure 4C**), but not DLS (**Figure 4D**), by one-way ANOVA in both limited (**Figure 4C**; n = 6 mice/group [19745 DMS genes], one-way ANOVA, main effect of DEG, F_(2, 19745)_ = 16.535, *p = 6.68e-08*; **Figure 4D**; n = 6 mice/group [19745 DLS genes], one-way ANOVA, main effect of DEG, F_(2, 11621)_ = 1.095, *p = 0.335*) and extended training (**Figures S11A–B**).

We next measured the relationship between CORT-induced differential H3K9ac enrichment and plasticity gene expression. Heatmaps were generated for plasticity DEGs from CORT versus vehicle comparisons that exhibited increased H3K9ac enrichment and increased differential gene expression, or decreased H3K9ac enrichment and decreased differential gene expression. CORT treatment, compared to vehicle groups, following limited training facilitated a strong female-specific relationship between decreased H3K9ac signal and decreased gene expression across DMS plasticity-related genes, while also exhibiting an increase in H3K9ac signal and increased gene expression across DLS plasticity-related genes (**Figures 4E–F**). Our data supported the hypothesis that gene expression differences correlate with H3K9ac levels in female DMS and DLS, suggesting sex- and region-specific differences in chromatin regulation either directly and/or indirectly due to chronic CORT and GR activation.

## Discussion

The present study found that CORT drives reward value insensitivity after limited operant conditioning in male and female mice, a time point when value-based flexible decision making would normally occur (**Figure S12**). We established that GR is necessary for the role of CORT in this context, using co-administration of a GR-antagonist. Our findings build on prior studies in male rodents, in which chronic stress or CORT induces a loss of flexible decision-making [12,17,18], manifesting as reward value insensitivity [24–26]. CORT-driven reward devaluation may reflect diminished positive valence of rewards [26,49–51]. Indeed, we found that chronic CORT reduced overall reward seeking while interfering with reward devaluation and accelerating the transition to inflexible behavior.

We sought to identify molecular mechanisms implicated in the transition to inflexible behavior. We found that the transition from limited to extended training was sufficient to reduce plasticity-related gene expression in the DMS. Remarkably, we found that CORT and limited training synergized to decrease plasticity-related gene expression in the DMS, and increase this in the DLS. Importantly, CORT accelerated the onset of plasticity gene expression changes, suggesting their functional relevance to the CORT-accelerated transition to inflexible behavior. GR directly regulates learning and memory consolidation [52], and acute GR-agonists produce a higher apoptotic response in rat DMS than DLS [53,54], implying striatal subregion-specific sensitivity to GR activation. The role of CORT in driving neuronal plasticity relevant to inflexible behavior is supported by previous studies in which stress or CORT increases dendritic density and complexity of DLS neurons [17,55] and DLS-dependent learning [56,57]. Alternatively, DMS loss-of-function enhances DLS-guided behavior acquisition [10,58,59], and stress reduces dendritic arbor complexity and plasticity in the DMS [17]. The current study offers subregion-specific gene-regulatory insights into the canonical transition between DMS- and DLS-guided behavior, in the presence or absence of CORT. One limitation is that our findings reflect the interaction effects of operant learning and chronic CORT on behavioral and molecular outputs.

Regarding mechanisms by which CORT regulates gene expression, we considered that chronic stress desensitizes neuronal GR and reduces downstream gene expression in the prefrontal cortex and hippocampus [60,61]. Our chronic CORT treatment also led to a global deenrichment of H3K9ac at DMS and DLS gene promoters following operant conditioning [48]. Our findings of CORT-reduced H3K9ac enrichment may be due to GR desensitization, which would be the first evidence for striatal-specific GR desensitization following chronic CORT. Alternatively, our chronic CORT may not desensitize GR in vivo, and downstream GR signaling from CORT treatment may be sufficient for the molecular changes observed. A limitation of our experimental design was only producing a chronic CORT state; future studies should consider the effects of acute CORT on operant and molecular outputs.

This study is the first to include both male and female mice in a study of CORT-driven behavioral inflexibility. We found very few DEGs between CORT-limited training and vehicle-extended training males, suggesting a single set of DEGs to promote inflexible behavior downstream of either extended or CORT-limited training. CORT may bring these gene expression patterns online earlier to produce inflexible decision-making phenotypes. Gene expression in both male and female mice showed subregion-specific activity patterns that correspond to loss of behavioral flexibility. Yet we found a high number of female-specific CORT-driven DEGs, SEs, and RI events, and correlated H3K9ac/DEG enrichment loci, suggesting that females use overlapping, but distinct, gene expression pathways downstream of extended or CORT-limited training to drive inflexible behavior. Other groups report similar sex-specific molecular phenomena concurrent with behavioral similarities. For example, CORT disrupts reward-seeking behaviors by decreasing dopamine reuptake in male DMS and reducing total dopamine levels in female DMS [62].

Beyond gene expression, we examined alternative splicing, which regulates >95% of expressed genes in neurons [41,42]. Critically, activity-dependent alternative splicing profiles are distinct from DEG profiles, yet few studies examine both modes of gene expression [43]. We found that plasticity genes are either differentially expressed or differentially spliced due to chronic CORT. SEs were the most commonly altered splicing event, which can influence transcript diversity and protein function [41]. Additionally, increased RIs in the DMS have the dual capacity to predispose transcripts for degradation, reducing expression [63] or increasing the proportion of nuclear-retained mRNA poised for export and translation under neuronal stimulation [64,65]. As such, repressed activity in the DMS in the transition to inflexible behavior may reflect both reduced expression and increased degradation of plasticity-related mRNA transcripts.

In sum, we discovered that DMS repression and DLS upregulation of synaptic plasticity gene expression accompany the natural and CORT-accelerated loss of behavioral flexibility in male and female mice. Our results support striatal subregion and sex-specific contributions to flexible decision-making via differential gene expression, alternative splicing, and permissive chromatin. Future endeavors should consider sex- and region-specific profiling of GR transcription factor binding or other chromatin modifications that may contribute to chronic stress-potentiated gene dysregulation and behavioral outcomes.

## Acknowledgments

Financial support was provided by NSF Graduate Research Fellowship Program (M.D.M., DGE-1845298), NIH T32 Training Grant (K.S.K., 5T32GM008216-37), NIH-NIMH Research Project Grant (E.A.H., R01-MH126027). We thank Dr. Shannon Gourley (Emory University) for critical review and revision of the written manuscript.

## Author contributions

Conceptualization and study design, M.D.M. and E.A.H. Animal husbandry and *in-vivo* experiments, M.D.M. Molecular experiments, M.D.M. RNAseq data analysis, K.S.K. and M.D.M. ChIPseq data analysis, S.Z. and M.D.M. Writing and editing, M.D.M. and E.A.H.

