## Supplemental Materials for "Corticosterone drives behavioral inflexibility via plasticity-related gene expression in the dorsal striatum"

### Supplemental Materials and Methods

#### Corticosterone and Mifepristone water administration

Corticosterone (CORT) (Sigma Aldrich, C2505, St. Louis, MO) and mifepristone (MIF) (Sigma-Aldrich, M8046) were dissolved in 100% molecular grade ethanol and then added to autoclaved mouse drinking water, such that the ultimate ethanol concentration was 1%. Vehicle-treated groups received 1% ethanol in their drinking water. Mice had ad libitum access to water, which was weighed and replaced with fresh solutions every two days. A CORT concentration of 50µg/mL was used for mice undergoing operant behavior. Water bottle weights and mouse body weights per cage were used to calculate the average CORT dose per cage of ~7mg/kg/day (**Figure 1B**). Mifepristone groups undergoing operant behavior received the same 50µg/mL of CORT plus 25µg/mL of MIF. The average MIF dose per cage was ~4mg/kg/day for the 25µg/mL groups (**Figure S4B**). Mice underwent two weeks of water administration before beginning behavior, and water administration continued throughout the duration of behavior.

#### Blood collection

Microcentrifuge collection tubes were kept on wet ice, and 100µL of 0.5 M EDTA was added to each sample tube prior to collection to avoid coagulation. Mice were allowed to freely move on cage tops, rather than restraining for blood collection, to avoid undue stress. 15-30µL blood samples were obtained by lateral tail vein laceration, aspirated with a 10µL pipette, and added to microcentrifuge sample tubes. Kwik Stop Styptic Powder (MiracleCorp, Dayton, OH) was applied with gauze to tail lacerations to clot bleeding and relieve pain after collection. Samples were centrifuged at 3000 rpm and 4 ºC for 15 min. Plasma was transferred to new microcentrifuge tubes and stored at -20 ºC. Blood samples were collected once a week for six weeks between 12:00PM and 1:30PM. Two night time points were selected between these weekly blood draws on night four and night ten and were conducted between 12:00AM and 1:30AM. Operant training and testing data for mice that experienced repeated blood collection have been omitted from our dataset due to the additional stress. Subsequent operant cohorts did not receive blood draws.

#### CORT ELISA

Blood plasma CORT was measured using a commercially available enzyme-linked immunosorbent assay (ELISA) kit (Abcam, AB108821, Cambridge, United Kingdom) according to the manufacturer's protocol. Briefly, plasma samples and a serial dilution of a stock CORT solution were added to a 96-well plate pre-coated with a CORT-specific antibody. Biotinylated CORT was then added, followed by washes. An avidin-biotin-peroxidase complex was added, and unbound conjugates were washed away. Finally, 3,3',5,5'-tetramethylbenzidine was catalyzed by hydrogen peroxide oxidoreductase to produce a blue color product that changed into yellow after adding an acidic stop solution. A microplate reader was then used to detect the absorbance of each well at 570 nm. The density of yellow coloration is inversely proportional to the amount of CORT captured by the plate, and the relative sample absorbance compared to the serial dilution can define the amount of CORT captured per sample. Either two-way or three-way analysis of the variance (ANOVA) tests were run with effect sizes reported as the standard omega squared ($\omega^2$) (% of total variation) for each main effect [1].

#### Operant response training

Training began after two weeks of water (Vehicle, CORT, or CORT+MIF) administration when mice were 11 weeks old. Mice were habituated to handling for three days prior to the initiation of training, and all training sessions were held between 7:30AM and 11:30AM. Sessions began by turning on the house light and ventilation fan within each sound-attenuating chamber (Med

Associates, MED-307W, Fairfax, VT). Each operant conditioning chamber was designated to one sex and water condition to avoid distracting scents. Mice underwent one session per day, during which pressing a lever initially resulted in a 20 mg sugar pellet reward outcome (Bio-Serv, F07595, Flemington, NJ). Mice progressed to the next training phase when at least 18 / 20 (90%) of possible reinforcers were earned within the two 40-minute sessions. The session would end when a subject earned 20 reinforcers or after 40-minutes. Food port entries, lever presses, reinforcers earned, and session times were recorded to compute food port entry rates and lever pressing rates.

Operant conditioning proceeded in a fashion similar to prior investigations [2–4]. After training mice according to a fixed ratio 1 (FR1) schedule of reinforcement, described above, mice graduated to a random interval 30s (RI-30s) schedule of reinforcement. Here, responses are not reinforced for randomized periods averaging 30s, after which, the next lever press is reinforced. RI schedules, with time, induce routinized habit-based response strategies because they uncouple the predictive relationship between actions and outcomes. Again, mice were required to accumulate 90% of available pellets to proceed to the next phase of the experiment. After three successful sessions of training according to the RI-30s schedule, mice graduated to an RI-60s schedule. Mice completed either three (limited training) or eight (extended training) sessions of RI-60s training, and the following day proceeded to the devaluation test. For molecular experiments, tissue was collected one hour after the mouse's final RI-60s training session.

#### Reward devaluation

Each mouse was placed individually in a clean empty cage without bedding, where it had free access to two grams of the same sugar pellets used as reinforcers for 1.5 hours. After freely consuming the reward, mice were transferred to their respective operant conditioning chambers.

Lever-pressing was monitored for 10 minutes in an unrewarded test. Since training responses (food port entries and lever presses) varied based on session timeout, response rates were calculated to inform reinforcer value. Data are expressed as % baseline, referring to the response rate generated during testing compared to the last two RI-60s training days, as previously described in the literature [5–10]. Raw lever pressing rates for the last two RI-60s sessions and the unrewarded test are also shown per mouse. Next, mice were returned to cages containing sucrose pellets for another 1.5 hours to repeat the consumption test. The purpose of the consumption test was to provide a measure for each subject's relative value for the sucrose pellet, with the expectation that after free consumption, the sucrose pellet would be devalued by the mice. Finally, mice were returned to their home cages. Pellet intake was measured by weighing the sugar pellets before and after each consumption session.

#### Tissue collection

Mice were euthanized 1 hour after their final RI-60s training session by cervical dislocation. The whole brain was immediately extracted and chilled for a few seconds in phosphate-buffered saline (PBS, pH 7.4) with cOmplete, EDTA-free Protease Inhibitor tablets (Roche, 4693159001, Manheim, Germany) on wet ice. The brain was then placed in a chilled slicer matrix, coronally sectioned at 1mm increments, and two slices (**Figure S5A**, Bregma +0.14mm A.P. and +1.18mm A.P., [11]), containing the regions of interest were placed in a sterile dish with chilled PBS + protease inhibitor. Bilateral 2mm-diameter biopsy punches were taken for each region of interest from two coronal slices (4 total punches per region per animal), placed in individual microcentrifuge tubes, and immediately flash frozen in a metal tube rack placed on dry ice. Samples were stored at -80 ºC. Given that the DMS and DLS have some degree of overlap in the central part between the substructures [12,13], samples potentially contained small amounts of the reciprocal subregion.

#### RNA and chromatin isolation for sequencing

The same tissue samples were processed for RNA and chromatin using single sample sequencing (S3EQ) [14–17]. Samples were submitted to Azenta Life Sciences (Plainfield, NJ) for sequencing, where RNA underwent additional quality control assays for RIN (8.5 to 9.9) and RNA concentration (28 - 98ng/µL). ChIP DNA underwent tape station (Azenta) and nanodrop (Azenta) analysis to ensure quality before sequencing.

#### Solution preparation for RNA and chromatin processing

The following solutions were prepared as previously described [14,15], with the addition of sodium butyrate (Sigma Aldrich, B5887) to prevent the removal of acetylation histone modifications.

Cell Lysis Buffer (10 mM Tris–HCl (pH 8.0), 10 mM NaCl, 3 mM MgCl2, 0.5% NP-40 in H2O, 10 mM sodium butyrate (Sigma Aldrich, B5887)),

BSA Blocking Buffer (0.5% Bovine serum albumin in 1X PBS),

Dilution Buffer (16.7 mM Tris–HCl (pH 8.0), 1.1% Triton-X 100, 0.01% SDS, 167 mM NaCl, 1.2 mM EDTA in H2O, 10 mM sodium butyrate),

Nuclear Lysis Buffer (50 mM Tris–HCl (pH 8.0), 5 mM EDTA, 1% SDS in H2O)

Wash Buffer 1 (20 mM Tris–HCl pH 8.0, 150 mM NaCl, 2 mM EDTA, 1% Triton X-100, 0.1% SDS in H2O, 10 mM sodium butyrate (Sigma Aldrich, B5887)),

Wash Buffer 2 (20 mM Tris–Cl pH 8.0, 500 mM NaCl, 2 mM EDTA, 1% Triton X-100, 0.1% SDS in H2O),

Wash Buffer 3 (250 mM LiCl, 10 mM Tris–HCl pH 8.0, 1% sodium deoxycholate, 1 mM EDTA, 1% IGEPAL CA-630 in H2O),

TE Buffer (10 mM Tris–HCl pH 8.0, 1 mM EDTA in H2O),

Elution Buffer (0.1 M NaHCO3, 1% SDS in H2O).

#### RNA extraction and purification

Tissue samples were homogenized in 185 µL Cell Lysis Buffer with a protease inhibitor (Roche, 4693159001) using a pellet pestle motor, and spun for five minutes (1000 xg, 4 ºC). The RNA-containing cytosolic supernatant and nuclei-containing pellet were separated and subjected to RNA extraction and purification or ChIP, respectively. RNA extraction and purification continued using the RNeasy Micro kit (Qiagen, 74004, Hilden, Germany). The cytosolic (RNA) fraction was mixed with 600 µL RLT buffer and 430µL 100% ethanol, followed by a 30-second spin (12,000 xg) in the RNeasy mini-spin columns. Columns were washed with 700µL of RW1 for a 30-second spin (12,000 xg), and then a DNAse solution was allowed to incubate on the column membranes for 15 min. Columns were spun twice for 30 seconds at 12,000 xg, first with 650 µL of RW1, then 500 µL of RPE. Then, two more 2-minute spins at 12,000 xg, first with 500 µL of RPE, then without liquid to dry the columns. 30 µL of RNase-free H2O was added to each column, and a 1-minute spin at 12,000 xg was used to elute RNA into fresh microcentrifuge tubes. RNA concentration was obtained using a Qubit 4 Fluorometer and RNA HS Assay Kit (Invitrogen, Eugene, OR). RNA was then flash-frozen on a metal tube rack placed on dry ice, then stored at -80 ºC.

#### Chromatin extraction and sheering

Chromatin processing continued using the nuclei pellet obtained from S3EQ. Pellets were resuspended in PBS and fixed with 11% formaldehyde for six minutes (350 rpm, 22 °C). 100 µL 1 M glycine was added, and samples were rocked for five minutes to stop cross-linking (500 rpm, 22 °C). Samples were centrifuged for five minutes (5500 × g, 4 °C), and the supernatant was discarded. The pellet was resuspended in 200 µL Nuclear Lysis Buffer, transferred to TPX tubes (Diagenode, C30010010, Denville, NJ), and incubated on ice for 10 minutes. Samples were then sonicated in a Bioruptor® (Diagenode) for three runs (high setting, 30 s on, 30 s off,

10 cycles). 10 µL of the 200 µL of chromatin was taken for quality control analysis. Dilution buffer was added to reach 1000 µL total. 10% input (relative to each IP) was collected from each sample for normalization in qChIP analysis. Samples and 10% input were then flash-frozen on a metal tube rack placed on dry ice, then stored at -80 ºC.

#### Chromatin immunoprecipitation (ChIP)

0.5X chromatin volume per sample of M280 Sheep anti-Rabbit Dynabeads (Invitrogen, 11204D) were washed three times in BSA blocking buffer. The beads were suspended in 1.5X bead volume of dilution buffer, and either a rabbit polyclonal H3K4me3 antibody (07–473, EMD Millipore, Burlington, MA) or a rabbit polyclonal H3K9ac antibody (C15410004, Diagenode) was added at a ratio of 1 µg antibody to 15 µL beads. The beads were then rotated for six hours (4 °C) to bind the antibody. The remaining diluted chromatin was thawed on wet ice, then combined with antibody-bound beads (1 µL bead-antibody slurry/1.6 µL chromatin) and placed on a rotator for 12 hours overnight (4 °C). Samples were then washed with 1 mL of ice-cold Wash Buffer 1, Wash Buffer 2, Wash Buffer 3, and TE Buffer using a DynaMag™-2 (Invitrogen, 12321D) to preserve beads between washes. Samples were rotated for five minutes during each wash (22 °C). Following washes, samples were rocked with 200 µL of elution buffer for 20 min (500 rpm, 22 °C), centrifuged for three minutes (14,000 xg), and placed back on the DynaMag™-2. The supernatant from each sample was transferred to fresh tubes. The 10 µL chromatin for QC and 10% input per sample were thawed on wet ice, and all volumes were brought to 200 µL using Elution Buffer.

#### DNA clean-up

The 10 µL chromatin for QC and 10% input per sample were thawed on wet ice, and all volumes were brought to 200 µL using Elution Buffer. 8 µL of 5 M NaCl and 2 µL of proteinase K 10 mg/mL) were added to each QC chromatin, 10% input, and ChIP-ed sample. All samples were incubated for 4 hours on a thermoblock (300 rpm, 65 ºC) for reverse crosslink and protein digestion. Proteinase K was heat-inactivated by incubating the samples for an extra 15 minutes (78 ºC), and then all DNA samples were purified using QIAmp Micro DNA kits (Qiagen, 56304). 200 µL of Buffer AL and 200 µL of 100% ethanol were mixed with each sample, then samples were transferred to QIAmp Mini Elute columns and spun for 1 minute (12,000 xg). Columns were spun twice for 1 minute at 12,000 xg, first with 500 µL of Buffer AW1, then 500 µL of Buffer AW2. 30 µL of ddH2O was added to each column, and a 1-minute spin at 12,000 xg was used to elute DNA into fresh microcentrifuge tubes. The QC chromatin was used to obtain DNA concentration from a Qubit 4 Fluorometer and DNA HS Assay Kit (Invitrogen), and DNA fragment size was verified at ~300bp with an Agilent 2100 bioanalyzer. Sample DNA and 10% inputs were then flash-frozen on a metal tube rack, placed on dry ice, then stored at -80 ºC.

#### Sequencing alignment and data analysis

Raw fastq files were trimmed to remove adapters, and read quality was measured using fastqc and multiqc. RNAseq reads were quality-controlled using FastQC [18] and MultiQC [19]. Then adapters were trimmed and aligned using kallisto to the mm39 genome. Aligned reads were then analyzed for differential expression using DESEQ2 [20] (version 1.38.3). All experimental groups were maintained for statistical analysis (i.e., n=3 per sex, brain region, water treatment, and training), and were within the standard number of samples and total comparisons for sequencing experiments [21,22]. The Benjamini-Hochberg correction for multiple hypotheses testing with a false discovery rate set to 0.05 was used to calculate adjusted p-values [23]. All sequencing data reported are adjusted p-values, and significance was attributed to adjusted p-values (FDR) for differential genes below 0.05. The R package clusterProfiler (version 4.14.3) was used to generate gene set enrichment analyses (GSEA), and adjusted p-values (q-values) were calculated using the Benjamini and Hochberg correction for multiple hypotheses testing with a false discovery rate set to 0.05 [24]. GSEA results were chosen by selecting the top five to ten gene set terms by adjusted p-value per group. Principal component analyses were conducted to verify replicate concordance based on experimental variables (**Figure S5B–E**). Several other R packages were used for data visualization. Raw sequencing files will be made available by request.

#### Splicing data analysis

To quantify differential alternative splicing as a consequence of corticosterone treatment, we utilized the rMATS [25] package to identify gene regions that were differentially included in the final transcript. Briefly, reads were aligned to the mm39 genome using the splice-aware aligner STAR using standard ENCODE parameters [26]. Aligned reads were then used to perform a paired analysis of differential splicing using rMATS. As reads were trimmed prior to alignment, the flag-variable-read-length was utilized to include trimmed reads as junction reads. The JCEC output files, which include both junction reads and on-target exonic reads to define PSI values, were used for all downstream visualization and analysis. Events were considered significant with an FDR < 0.05 and |DPSI| > 5%. These cutoffs have been used by the field to classify events that can be robustly validated by biochemical methods.

#### ChIP-seq data analysis

Paired-end raw reads were mapped to the mm39 reference genome using bowtie2(v2.1.0) [27] with parameters:-q--local--very-sensitive--no-mixed--no-unal--dovetail--phred33. Properly and uniquely aligned reads were selected using samtools view function (v1.9) [28] with parameters:-bS-q 20-f 0x2. Then, picard (v2.23.4) [29] was used to remove duplicates from filtered read pairs.Reads overlapping with blacklist regions [30] were further removed using bedtools intersect function (v2.29.2) [31]. Peaks of each H3K9ac/GR ChIP-seq library were called by MACS2 [32] with corresponding input as control with parameters:-f BAMPE-g mm-q 0.01. Additional one-way ANOVAs between H3K9ac enrichment and differentially expressed genes were performed using the R package, “rstatix” [33].

#### Data availability

The datasets used and/or analysed during the current study are available from the corresponding author on reasonable request.

#### Supplemental Figures S1 – S12

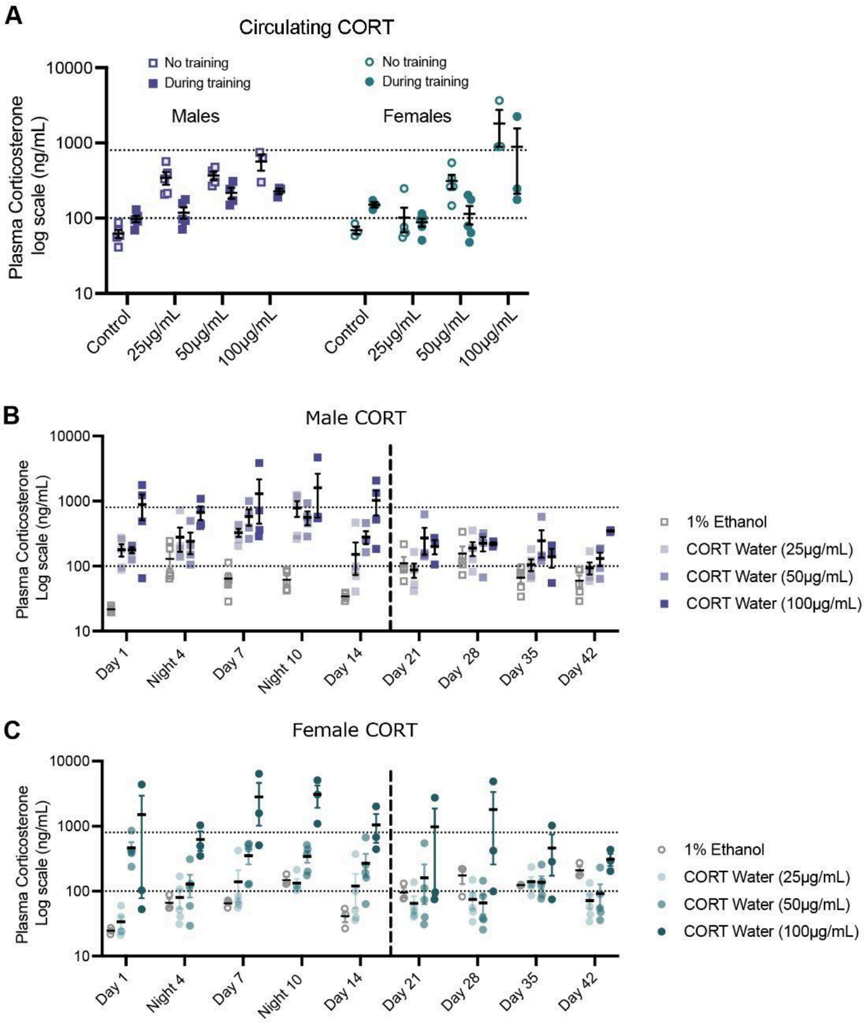

**Figure S1: CORT administration produces elevated plasma CORT. A** Circulating CORT increased with increasing chronic CORT dose from 25µg/mL to 100µg/mL [34–36], and CORT (50µg/mL) produced the most consistently elevated levels of CORT in both males and females [n = 3-5 mice/group, three-way ANOVA, main effect of Training, $F_{(1, 50)} = 4.426$, $p = 0.0404$; main effect of Sex, $F_{(1, 50)} = 3.483$, $p = 0.0679$; main effect of CORT Dose, $F_{(3, 50)} = 10.04$, $p < 0.0001$]. Physiological cutoffs for chronic stress-like levels were set between 100 – 800ng/mL [37,38]. Data shown as log10 to account for the ten-fold change in circulating CORT between controls and low or moderate dose, and a hundred-fold change between controls and the high dose group. **B, C** Male [n = 3-5 mice/group, two-way ANOVA, main effect of Time, $F_{(8, 104)} = 6.171$, $p < 0.0001$; main effect of CORT Dose, $F_{(3, 13)} = 14.15$, $p = 0.0002$] and female [n = 3-5 mice/group, two-way ANOVA, main effect of Time, $F_{(8, 96)} = 4.128$, $p = 0.0003$; main effect of CORT Dose, $F_{(3, 12)} = 3.859$, $p = 0.0382$] plasma CORT levels increased with increasing CORT dose. There was no difference in plasma CORT between day and night. The onset of operant conditioning (vertical line) transiently increased plasma CORT for vehicle groups, but not for CORT administration groups. See **Table S1** for detailed statistics.

| Figure | Metric | ANOVA comparison | F(DFn, DFd) | p value | ω^2 |
|---|---|---|---|---|---|
| **Fig. S1A** | Circulating CORT | Training | F(1, 50) = 4.426 | 0.0404 | 4.082 |
| | | Sex | F(1, 50) = 3.483 | 0.0679 | 3.212 |
| | | CORT Dose | F(3, 50) = 10.04 | < 0.0001 | 27.79 |
| | | Training x Sex | F(1, 50) = 0.2140 | 0.6456 | 0.1974 |
| | | Training x CORT Dose | F(3, 50) = 1.700 | 0.1789 | 4.704 |
| | | Sex x CORT Dose | F(3, 50) = 5.304 | 0.003 | 14.67 |
| | | Training x Sex x CORT Dose | F(3, 50) = 0.6328 | 0.5973 | 1.751 |
| **Fig. S1B** | Male CORT | Time | F(8, 104) = 6.171 | < 0.0001 | 16.67 |
| | | CORT Dose | F(3, 13) = 14.15 | 0.0002 | 21.31 |
| | | Time x CORT Dose | F(24, 104) = 2.502 | 0.0008 | 20.27 |
| **Fig. S1C** | Female CORT | Time | F(8, 96) = 4.128 | 0.0003 | 7.323 |
| | | CORT Dose | F(3, 12) = 3.859 | 0.0382 | 28.66 |
| | | Time x CORT Dose | F(24, 96) = 2.888 | 0.0001 | 15.37 |

**Table S1: Detailed statistics for Figure S1.** Either two-way or three-way analysis of the variance (ANOVA) tests were run with effect sizes reported as the standard omega squared ($\omega^2$) (% of total variation) for each main effect [1]. Corresponding figures, metrics, ANOVA comparisons, F statistics, and p values are shown.

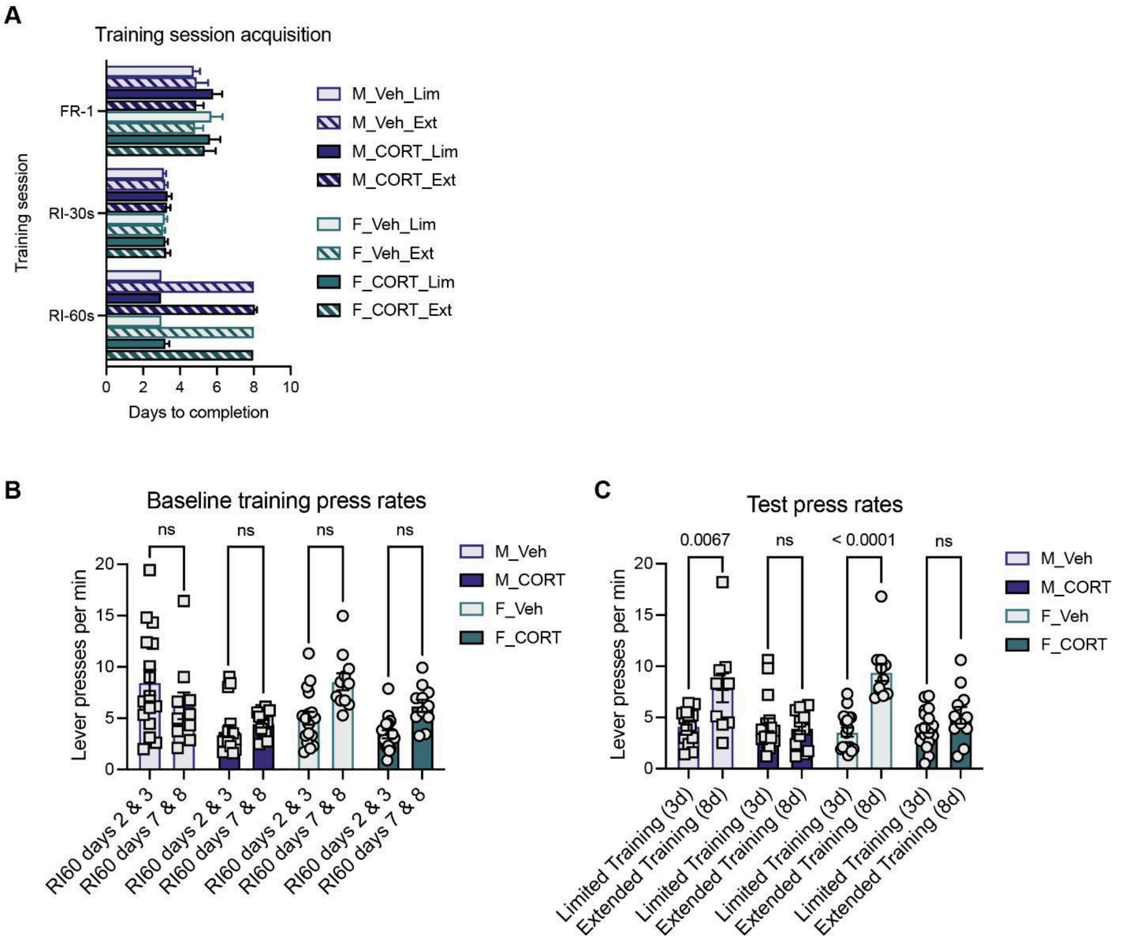

**Figure S2: CORT reduces operant task engagement and increases tested behavioral inflexibility.** **A** There was no CORT or sex difference in the number of days it took for mice to progress to the next training session within limited [n = 16-17 mice/group, three-way ANOVA, main effect of Training Session, $F_{(2, 171)} = 67.35$, *p < 0.0001*; main effect of CORT, $F_{(1, 171)} = 1.074$, *p = 0.3016*; main effect of Sex, $F_{(1, 171)} = 0.2843$, *p = 0.5946*] or extended groups [n = 10-12 mice/group, three-way ANOVA, main effect of Training Session, $F_{(2, 120)} = 257.1$, *p < 0.0001*; main effect of CORT, $F_{(1, 120)} = 0.8404$, *p = 0.3611*; main effect of Sex, $F_{(1, 120)} = 0.01145$, *p = 0.915*]. **B** CORT reduced training press rates [n = 10-17 mice/group, three-way ANOVA, main effect of Training Duration, $F_{(1, 103)} = 4.353$, *p = 0.0394*; main effect of CORT, $F_{(1, 103)} = 20.49$, *p < 0.0001*; main effect of Sex, $F_{(1, 103)} = 0.01839$, *p = 0.8924*], however there was no significant difference in baseline training press rates between limited and extended groups [n = 10-17 mice/group, three-way ANOVA followed by Tukey post-hoc tests, Males Vehicle: Limited v Extended, DF = 103, *p = 0.5272*; Males CORT: Limited v Extended, DF = 103, *p = 0.9977*; Females Vehicle: Limited v Extended, DF = 103, *p = 0.1967*; Females CORT: Limited v Extended, DF = 103, *p = 0.2162*]. **C** Test lever pressing rates significantly differed for vehicle but not CORT groups, and did not differ by sex [n = 10-17 mice/group, three-way ANOVA followed by Tukey post-hoc tests, main effect of Training Duration, $F_{(1, 103)} = 29.85$, *p < 0.0001*;

main effect of Sex, $F_{(1, 103)} = 0.9448$, *p = 0.3333*; main effect of CORT, $F_{(1, 103)} = 15.77$, *p = 0.0001*; Males Vehicle: Limited v Extended, DF = 103, *p = 0.0067*; Males CORT: Limited v Extended, DF = 103, *p > 0.9999*; Females Vehicle: Limited v Extended, DF = 103, *p < 0.0001*; Females CORT: Limited v Extended, DF = 103, *p = 0.8874*]. See **Table S2** for detailed statistics.

| Figure | Metric | ANOVA comparison | F(DFn, DFd) | p value |
|---|---|---|---|---|
| **Fig. S2A** | Training session acquisition - limited training | Training Session | F (2, 171) = 67.35 | < 0.0001 |
| | | CORT | F (1, 171) = 1.074 | 0.3016 |
| | | Sex | F (1, 171) = 0.2843 | 0.5946 |
| | | Training Session x CORT | F (2, 171) = 0.2145 | 0.8072 |
| | | Training Session x Sex | F (2, 171) = 0.2479 | 0.7808 |
| | | CORT x Sex | F (1, 171) = 0.4017 | 0.5271 |
| | | Training Session x CORT x Sex | F (2, 171) = 0.7235 | 0.4865 |
| **Fig. S2A** | Training session acquisition - extended training | Training Session | F (2, 120) = 257.1 | < 0.0001 |
| | | CORT | F (1, 120) = 0.8404 | 0.3611 |
| | | Sex | F (1, 120) = 0.01145 | 0.915 |
| | | Training Session x CORT | F (2, 120) = 0.1566 | 0.8552 |
| | | Training Session x Sex | F (2, 120) = 0.2253 | 0.7986 |
| | | CORT x Sex | F (1, 120) = 0.2222 | 0.6382 |
| | | Training Session x CORT x Sex | F (2, 120) = 0.2836 | 0.7536 |
| **Fig. S2B** | Baseline training press rates | Training duration | F (1, 103) = 4.353 | 0.0394 |
| | | Sex | F (1, 103) = 0.01839 | 0.8924 |
| | | CORT | F (1, 103) = 20.49 | < 0.0001 |
| | | Training duration x Sex | F (1, 103) = 11.74 | 0.0009 |
| | | Training duration x CORT | F (1, 103) = 0.9262 | 0.3381 |

| | | | | |
|---|---|---|---|---|
| | | Sex x CORT | F (1, 103) = 1.273 | 0.2618 |
| | | Training duration x Sex x CORT | F (1, 103) = 2.688 | |
| **Fig. S2B** | Baseline training press rates | Males Vehicle: RI60s D2&3 v D7&8 (Tukey post-hoc) | DF = 103 | 0.5272 |
| | | Males CORT: RI60s D2&3 v D7&8 (Tukey post-hoc) | DF = 103 | 0.9977 |
| | | Females Vehicle: RI60s D2&3 v D7&8 (Tukey post-hoc) | DF = 103 | 0.1967 |
| | | Females CORT: RI60s D2&3 v D7&8 (Tukey post-hoc) | DF = 103 | 0.2162 |
| **Fig. S2C** | Test press rates | Training duration | F (1, 103) = 29.85 | < 0.0001 |
| | | Sex | F (1, 103) = 0.9448 | 0.3333 |
| | | CORT | F (1, 103) = 15.77 | 0.0001 |
| | | Training duration x Sex | F (1, 103) = 3.746 | 0.0557 |
| | | Training duration x CORT | F (1, 103) = 20.37 | < 0.0001 |
| | | Sex x CORT | F (1, 103) = 3.484e-005 | 0.9953 |
| | | Training duration x Sex x CORT | F (1, 103) = 0.06112 | 0.8052 |
| **Fig. S2C** | Test press rates | Males Vehicle: RI60s D2&3 v D7&8 (Tukey post-hoc) | DF = 103 | 0.0067 |
| | | Males CORT: RI60s D2&3 v D7&8 (Tukey post-hoc) | DF = 103 | > 0.9999 |
| | | Females Vehicle: RI60s D2&3 v D7&8 (Tukey post-hoc) | DF = 103 | < 0.0001 |
| | | Females CORT: RI60s D2&3 v D7&8 (Tukey post-hoc) | DF = 103 | 0.8874 |

**Table S2: Detailed statistics for Figure S2.** Three-way analysis of the variance (ANOVA) tests were run with a Tukey post-hoc to correct for multiple comparisons. Corresponding figures, metrics, ANOVA comparisons, F statistics, and p values are shown.

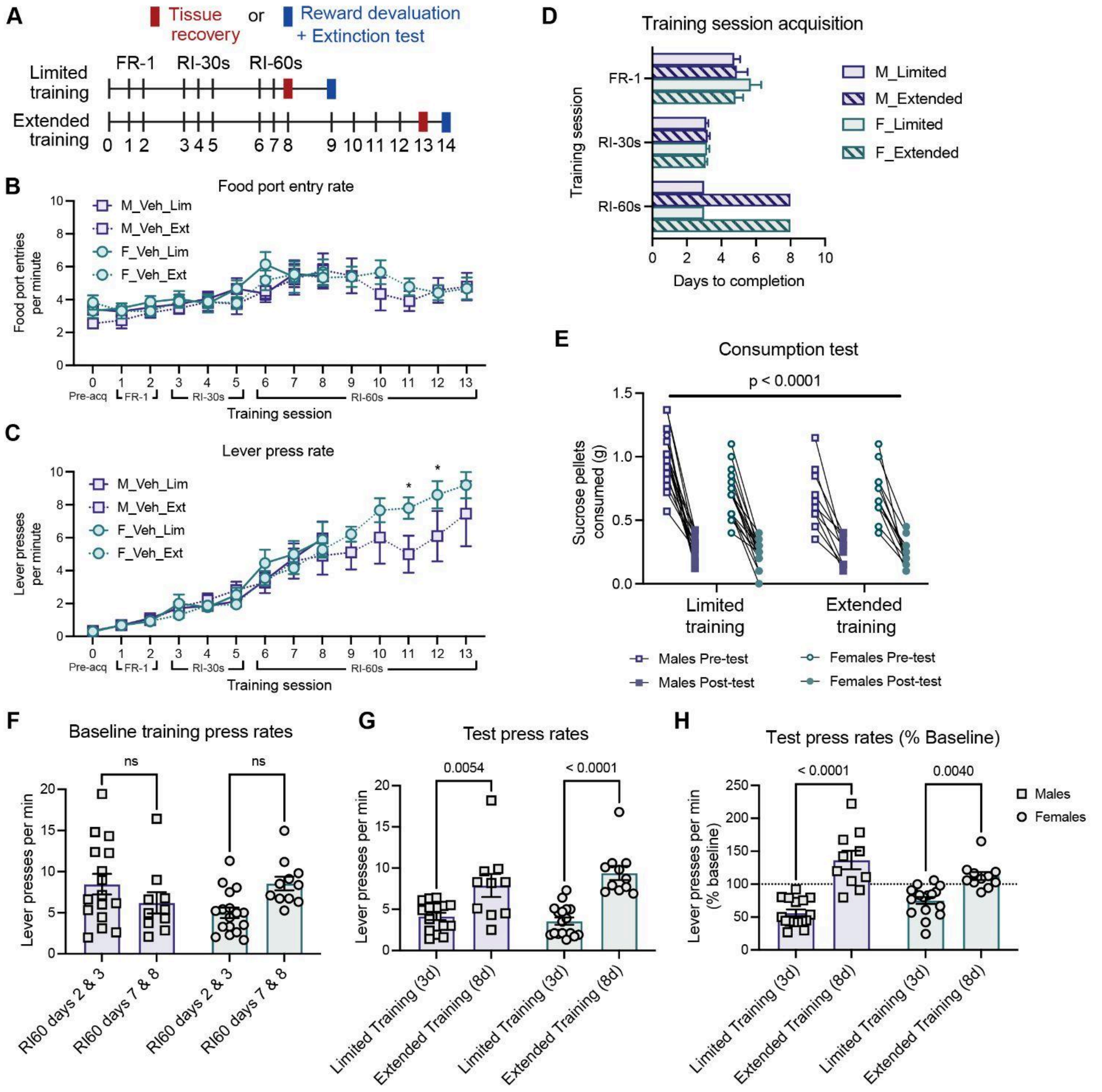

**Figure S3: An extended number of operant training sessions promotes inflexible behavior in male and female mice. A** Behavioral timeline of limited or extended operant conditioning. All mice underwent two days of FR-1 training, three days of RI-30s training, then either three days of RI-60s training for limited training groups, or eight days of RI-60s training for extended training groups. Mice progressed to a reward devaluation and testing the day after their final RI-60s training session. **B, C** The average food port entry rate per minute or lever pressing rate per minute ± SEM is shown across each day of the operant learning task. The average food port entry rate [n = 10-17 mice/group, two-way ANOVA, main effect of Training Day, $F_{(13, 508)}$ = 11.69, *p < 0.0001*; main effect of Sex, $F_{(1, 52)}$ = 0.2301, *p = 0.6335*] and lever pressing rate [n = 10-17 mice/group, two-way ANOVA, main effect of Training Day, $F_{(13, 508)}$ = 73.3, *p < 0.0001*; main effect of Sex, $F_{(1, 52)}$ = 0.09666, *p = 0.7571*] for males and females were not different during behavioral acquisition. **D** There was no sex difference in the number of days it took for mice to progress to the next training session [n = 10-17 mice/group, three-way ANOVA, main effect of Training Session, $F_{(2, 100)}$ = 60.68, *p < 0.0001*; main effect of Training Duration, $F_{(1, 50)}$ = 73.17, *p < 0.0001*; main effect of Sex, $F_{(1, 50)}$ = 0.5698, *p = 0.4539*]. **E** All mice reduced free consumption of the sucrose reward following testing compared to their pre-test consumption during sensory-specific satiety devaluation [n = 10-17 mice/group, two-way ANOVA, main effect of Pre v Post Consumption Test, $F_{(1, 52)}$ = 286.3, *p < 0.0001*]. **F** There was no difference in baseline training press rates based on training duration (limited or extended) or sex [n = 10-17 mice/group, two-way ANOVA, main effect of Training Duration, $F_{(1, 50)}$ = 0.3759, *p = 0.5426*; main effect of Sex, $F_{(1, 50)}$ = 0.2932, *p = 0.5906*]. **G** Test lever pressing rates significantly differed based on training duration, but not by sex [n = 10-17 mice/group, two-way ANOVA, main effect of Training Duration, $F_{(1, 50)}$ = 41.68, *p < 0.0001*; main effect of Sex, $F_{(1, 50)}$ = 0.4005, *p = 0.5297*]. **H** Mice tested following limited training reduced their lever pressing rate below their individual baselines, while mice undergoing extended training increased lever pressing rates during testing [n = 10-17 mice/group, two-way ANOVA followed by Tukey post-hoc tests, main effect of Training Duration, $F_{(1, 50)}$ = 58.31, *p < 0.0001*; Males: Limited v Extended, DF = 50, *p < 0.0001*; Females: Limited v Extended, DF = 50, *p = 0.0040*]. Individual baselines were defined by the lever-pressing rate on the two previous days of RI-60s training. See **Table S3** for detailed statistics.

| Figure | Metric | ANOVA comparison | F(DFn, DFd) | p value |
|---|---|---|---|---|
| **Fig. S3B** | Food port entry rate - limited training | Training Day | F (8, 208) = 8.421 | < 0.0001 |
| | | Sex | F (1, 26) = 0.1348 | 0.7165 |
| | | Training Day x Sex | F (8, 208) = 0.8222 | 0.5838 |
| **Fig. S3B** | Food port entry rate - limited training | M_v_F D0 (Šídák post-hoc) | DF = 234 | 0.7215 |
| | | M_v_F D1 (Šídák post-hoc) | DF = 234 | 0.8903 |
| | | M_v_F D2 (Šídák post-hoc) | DF = 234 | 0.8732 |
| | | M_v_F D3 (Šídák post-hoc) | DF = 234 | 0.5747 |
| | | M_v_F D4 (Šídák post-hoc) | DF = 234 | 0.3497 |

| | | | | |
|---|---|---|---|---|
| | | M_v_F D5 (Šídák post-hoc) | DF = 234 | 0.3043 |
| | | M_v_F D6 (Šídák post-hoc) | DF = 234 | 0.2092 |
| | | M_v_F D7 (Šídák post-hoc) | DF = 234 | 0.5445 |
| | | M_v_F D8 (Šídák post-hoc) | DF = 234 | 0.7409 |
| **Fig. S3B** | Food port entry rate - extended training | Training Day | F (13, 244) = 7.402 | < 0.0001 |
| | | Sex | F (1, 19) = 0.3558 | 0.5579 |
| | | Training Day x Sex | F (13, 244) = 0.8580 | 0.5982 |
| **Fig. S3B** | Food port entry rate - extended training | M_v_F D0 (Šídák post-hoc) | DF = 263 | 0.0611 |
| | | M_v_F D1 (Šídák post-hoc) | DF = 263 | 0.5183 |
| | | M_v_F D2 (Šídák post-hoc) | DF = 263 | 0.9256 |
| | | M_v_F D3 (Šídák post-hoc) | DF = 263 | 0.6335 |
| | | M_v_F D4 (Šídák post-hoc) | DF = 263 | 0.9881 |
| | | M_v_F D5 (Šídák post-hoc) | DF = 263 | 0.9519 |
| | | M_v_F D6 (Šídák post-hoc) | DF = 263 | 0.4654 |
| | | M_v_F D7 (Šídák post-hoc) | DF = 263 | 0.7212 |
| | | M_v_F D8 (Šídák post-hoc) | DF = 263 | 0.5958 |
| | | M_v_F D9 (Šídák post-hoc) | DF = 263 | 0.9446 |
| | | M_v_F D10 (Šídák post-hoc) | DF = 263 | 0.1302 |
| | | M_v_F D11 (Šídák post-hoc) | DF = 263 | 0.3246 |
| | | M_v_F D12 (Šídák post-hoc) | DF = 263 | 0.8652 |
| | | M_v_F D13 (Šídák post-hoc) | DF = 263 | 0.8968 |
| **Fig. S3C** | Lever pressing rate - limited training | Training Day | F (8, 200) = 45.76 | < 0.0001 |
| | | Sex | F (1, 25) = 0.01602 | 0.9003 |
| | | Training Day x Sex | F (8, 200) = 0.3791 | 0.9309 |

| | | | | |
|---|---|---|---|---|
| **Fig. S3C** | Lever pressing rate - limited training | M_v_F D0 (Šídák post-hoc) | DF = 234 | 0.9958 |
| | | M_v_F D1 (Šídák post-hoc) | DF = 234 | 0.9688 |
| | | M_v_F D2 (Šídák post-hoc) | DF = 234 | 0.9239 |
| | | M_v_F D3 (Šídák post-hoc) | DF = 234 | 0.9894 |
| | | M_v_F D4 (Šídák post-hoc) | DF = 234 | 0.9126 |
| | | M_v_F D5 (Šídák post-hoc) | DF = 234 | 0.7847 |
| | | M_v_F D6 (Šídák post-hoc) | DF = 234 | 0.5024 |
| | | M_v_F D7 (Šídák post-hoc) | DF = 234 | 0.8052 |
| | | M_v_F D8 (Šídák post-hoc) | DF = 234 | 0.2121 |
| **Fig. S3C** | Lever pressing rate - limited training | Training Day | F (13, 244) = 45.36 | < 0.0001 |
| | | Sex | F (1, 19) = 0.5906 | 0.4516 |
| | | Training Day x Sex | F (13, 244) = 2.138 | 0.0128 |
| **Fig. S3C** | Lever pressing rate - limited training | M_v_F D0 (Šídák post-hoc) | DF = 263 | 0.9348 |
| | | M_v_F D1 (Šídák post-hoc) | DF = 263 | 0.9923 |
| | | M_v_F D2 (Šídák post-hoc) | DF = 263 | 0.9592 |
| | | M_v_F D3 (Šídák post-hoc) | DF = 263 | 0.6584 |
| | | M_v_F D4 (Šídák post-hoc) | DF = 263 | 0.7693 |
| | | M_v_F D5 (Šídák post-hoc) | DF = 263 | 0.4289 |
| | | M_v_F D6 (Šídák post-hoc) | DF = 263 | 0.8062 |
| | | M_v_F D7 (Šídák post-hoc) | DF = 263 | 0.6947 |
| | | M_v_F D8 (Šídák post-hoc) | DF = 263 | 0.7369 |
| | | M_v_F D9 (Šídák post-hoc) | DF = 263 | 0.3168 |
| | | M_v_F D10 (Šídák post-hoc) | DF = 263 | 0.131 |
| | | M_v_F D11 (Šídák post-hoc) | DF = 263 | 0.0107 |
| | | M_v_F D12 (Šídák post-hoc) | DF = 263 | 0.022 |
| | | M_v_F D13 (Šídák post-hoc) | DF = 263 | 0.1132 |

| | | | | |
|---|---|---|---|---|
| **Fig. S3D** | Training session acquisition | Training Session | F (2, 100) = 60.68 | < 0.0001 |
| | | Training Duration | F (1, 50) = 73.17 | < 0.0001 |
| | | Sex | F (1, 50) = 0.5698 | 0.4539 |
| | | Training Session x Training Duration | F (2, 100) = 87.79 | < 0.0001 |
| | | Training Session x Sex | F (2, 100) = 0.6646 | 0.5168 |
| | | Training Duration x Sex | F (1, 50) = 1.227 | 0.2732 |
| | | Training Session x Training Duration x Sex | F (2, 100) = 0.7607 | 0.47 |
| **Fig. S3E** | Consumption test | Pre v Post | F(1, 52) = 286.3 | < 0.0001 |
| | | Sex | F(1, 52) = 5.505 | 0.0228 |
| | | Pre v Post x Sex | F(1, 52) = 4.431 | 0.0401 |
| **Fig. S3F** | Baseline training press rates | Training Duration | F (1, 50) = 0.3759 | 0.5426 |
| | | Sex | F (1, 50) = 0.2932 | 0.5906 |
| | | Training Duration x Sex | F (1, 50) = 7.634 | 0.008 |
| **Fig. S3F** | Baseline training press rates | Males: RI60s D2&3 v D7&8 (Tukey post-hoc) | DF = 50 | 0.451 |
| | | Females: RI60s D2&3 v D7&8 (Tukey post-hoc) | DF = 50 | 0.083 |
| | | RI60s D2&3: Males v Females (Tukey post-hoc) | DF = 50 | 0.0509 |
| | | RI60s D7&8: Males v Females (Tukey post-hoc) | DF = 50 | 0.4928 |
| **Fig. S3G** | Test press rates | Training Duration | F (1, 50) = 41.68 | < 0.0001 |
| | | Sex | F (1, 50) = 0.4005 | 0.5297 |

| | | | | |
|---|---|---|---|---|
| | | Training Duration x Sex | F (1, 50) = 1.995 | 0.164 |
| **Fig. S3G** | Baseline training press rates | Males: Limited v Extended (Tukey post-hoc) | DF = 50 | 0.0054 |
| | | Females: Limited v Extended (Tukey post-hoc) | DF = 50 | < 0.0001 |
| | | Limited: Males v Females (Tukey post-hoc) | DF = 50 | 0.9235 |
| | | Extended: Males v Females (Tukey post-hoc) | DF = 50 | 0.5624 |
| **Fig. S3H** | Test press rates | Training Duration | F (1, 50) = 62.36 | < 0.0001 |
| | | Sex | F (1, 50) = 0.1343 | 0.7156 |
| | | Training Duration x Sex | F (1, 50) = 8.490 | 0.0053 |
| **Fig. S3H** | Test press rate (% Baseline) | Males: Limited v Extended (Tukey post-hoc) | DF = 50 | < 0.0001 |
| | | Females: Limited v Extended (Tukey post-hoc) | DF = 50 | 0.004 |
| | | Limited: Male v Female (Tukey post-hoc) | DF = 50 | 0.1862 |
| | | Extended: Male v Female (Tukey post-hoc) | DF = 50 | 0.168 |

**Table S3: Detailed statistics for Figure S3.** Either two-way or three-way analysis of the variance (ANOVA) tests were run with either a Šidák or Tukey post-hoc to correct for multiple comparisons. Corresponding figures, metrics, ANOVA comparisons, F statistics, and p values are shown.

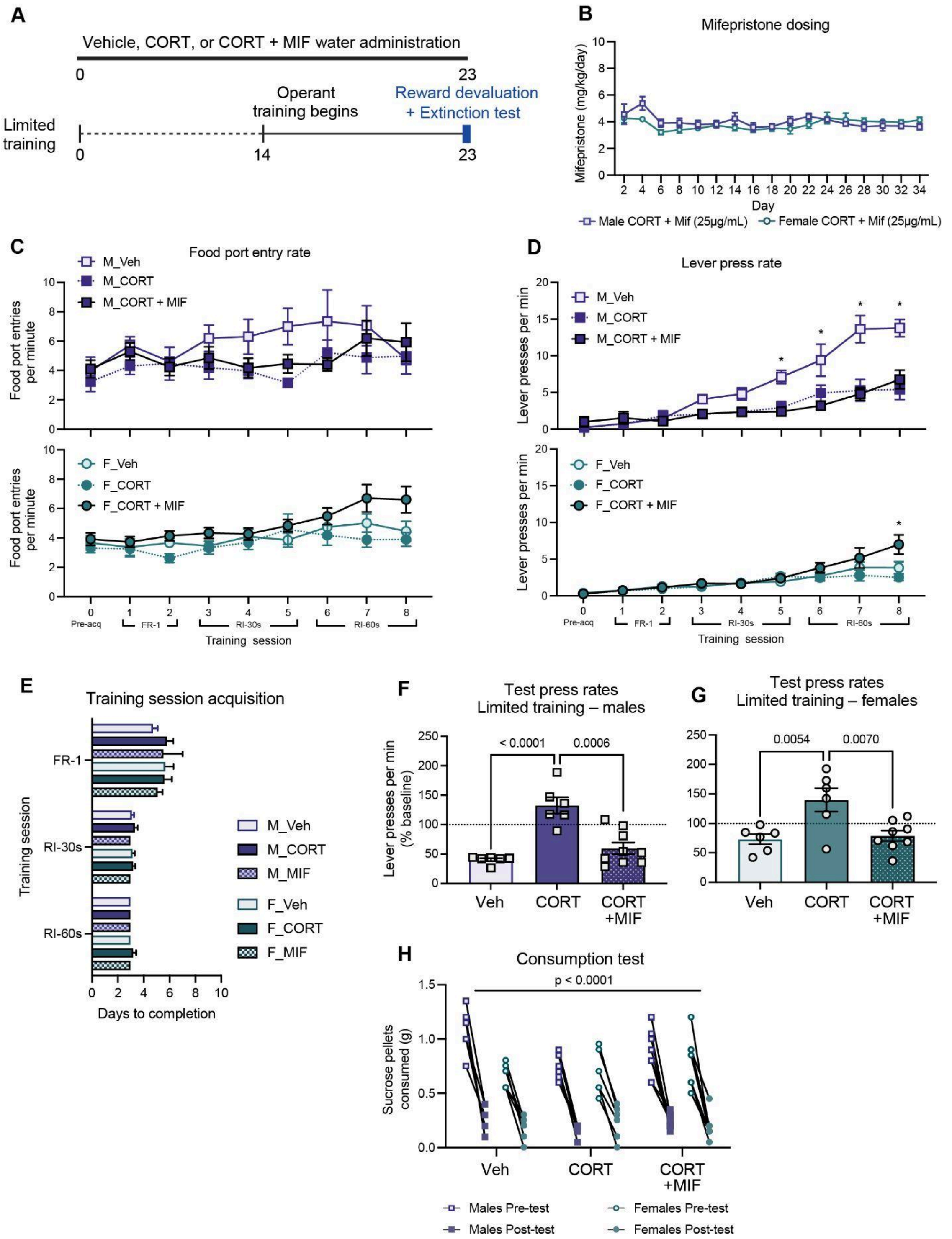

**Figure S4: CORT-accelerated loss of behavioral flexibility is attributable to GR binding.** **A** Behavioral timeline where vehicle, CORT, or CORT + MIF water administration began 14 days before training and lasted throughout the duration of limited operant training and testing. **B** There was no sex difference in MIF dosing when adjusting for body weight and water intake per cage [n = 3 cages/group, two-way ANOVA, main effect of Time, $F_{(16, 64)} = 4.619$, *p < 0.0001*; main effect of Sex, $F_{(1, 4)} = 0.397$, *p = 0.5628*]. **C, D** The average food port entry rate per minute or lever pressing rate per minute ± SEM by sex (male or female) and treatment (vehicle, CORT, or CORT + MIF) is shown across each day of the operant conditioning task. CORT had no effect on food port entries [n = 6-9 mice/group, three-way ANOVA, main effect of Training Day, $F_{(8, 208)} = 5.122$, *p < 0.0001*; main effect of Sex, $F_{(1, 26)} = 3.78$, *p = 0.0628*; main effect of Treatment, $F_{(1, 26)} = 0.1190$, *p = 0.7329*]. Treatment with CORT or CORT + MIF decreased the male lever pressing rates [n = 6-9 mice/group, three-way ANOVA, main effect of Training Day, $F_{(8, 208)} = 67.32$, *p < 0.0001*; main effect of Sex, $F_{(1, 26)} = 18.38$, *p = 0.0002*; main effect of Treatment, $F_{(1, 26)} = 7.004$, *p = 0.0136*]. All mice exhibited an increase in lever pressing as the random interval for training increased. **E** There was no treatment or sex difference in the number of days it took for mice to progress to the next training session [n = 6-9 mice/group, three-way ANOVA, main effect of Training Session, $F_{(2, 78)} = 18.14$, *p < 0.0001*; main effect of Treatment, $F_{(1, 78)} = 0.02273$, *p = 0.8806*; main effect of Sex, $F_{(1, 78)} = 0.7299$, *p = 0.3955*]. **F, G** In an unrewarded lever pressing test following sensory-specific satiety devaluation, vehicle and CORT + MIF groups attenuate lever pressing while CORT groups do not attenuate lever pressing and display an inflexible lever pressing response [n = 6-9 mice/group, two-way ANOVA followed by Tukey post-hoc tests, main effect of Treatment, $F_{(1, 35)} = 4.3342$, *p < 0.0001*; main effect of Sex, $F_{(2, 35)} = 25.48$, *p = 0.0446*; VehMales v CORTMales, DF = 35, *p < 0.0001*; VehMales v MIFMales, DF = 35, *p = 0.8193*; CORTMales v MIFMales, DF = 35, *p = 0.0006*; VehFemales v CORTFemales, DF = 35, *p = 0.0054*; VehFemales v MIFFemales, DF = 35, *p = 0.9991*; CORTFemales v MIFFemales, DF = 35, *p = 0.007*]. Lever presses per minute from the test are reported as the percent of an individual's baseline from the average of the individual's last two completed training sessions. **H** All mice reduced free consumption of the sucrose reward following the test compared to their pre-test consumption during sensory-specific satiety devaluation [n = 6-9 mice/group, three-way ANOVA, main effect of Pre v Post Consumption Test, $F_{(1, 26)} = 224.9$, *p < 0.0001*; main effect of Sex, $F_{(1, 26)} = 12.87$, *p = 0.0014*; main effect of Treatment, $F_{(1, 26)} = 0.3385$, *p = 0.5657*]. See **Table S4** for detailed statistics.

| Figure | Metric | ANOVA comparison | F(DFn, DFd) | p value |
|---|---|---|---|---|
| **Fig. S4B** | Mifepristone dosing | Time | F(16, 64) = 4.619 | < 0.0001 |
| | | Sex | F(1, 4) = 0.397 | 0.5628 |
| | | Time x Sex | F(16, 64) = 2.404 | 0.0069 |
| **Fig. S4C** | Food port entry rate | Training Day | F(8, 208) = 5.122 | < 0.0001 |
| | | Sex | F(1, 26) = 3.78 | 0.0628 |
| | | Treatment | F(1, 26) = 0.1190 | 0.7329 |

| | | | | |
|---|---|---|---|---|
| | | Training day x Sex | F(8, 208) = 1.442 | 0.1806 |
| | | Training Day x Treatment | F(8, 208) = 2.119 | 0.0354 |
| | | Sex x Treatment | F(1, 26) = 4.079 | 0.0538 |
| | | Training Day x Sex x Treatment | F(8, 208) = 1.45 | 0.1774 |
| **Fig. S4C** | Food port entry rate | Males: Vehicle v MIF D0 (Šídák post-hoc) | DF = 234 | > 0.9999 |
| | | Males: Vehicle v MIF D1 (Šídák post-hoc) | DF = 234 | > 0.9999 |
| | | Males: Vehicle v MIF D2 (Šídák post-hoc) | DF = 234 | > 0.9999 |
| | | Males: Vehicle v MIF D3 (Šídák post-hoc) | DF = 234 | > 0.9999 |
| | | Males: Vehicle v MIF D4 (Šídák post-hoc) | DF = 234 | > 0.9999 |
| | | Males: Vehicle v MIF D5 (Šídák post-hoc) | DF = 234 | 0.2148 |
| | | Males: Vehicle v MIF D6 (Šídák post-hoc) | DF = 234 | 0.9983 |
| | | Males: Vehicle v MIF D7 (Šídák post-hoc) | DF = 234 | > 0.9999 |
| | | Males: Vehicle v MIF D8 (Šídák post-hoc) | DF = 234 | > 0.9999 |
| **Fig. S4C** | Food port entry rate | Females: Vehicle v MIF D0 (Šídák post-hoc) | DF = 234 | > 0.9999 |
| | | Females: Vehicle v MIF D1 (Šídák post-hoc) | DF = 234 | > 0.9999 |
| | | Females: Vehicle v MIF D2 (Šídák post-hoc) | DF = 234 | > 0.9999 |
| | | Females: Vehicle v MIF D3 (Šídák post-hoc) | DF = 234 | > 0.9999 |
| | | Females: Vehicle v MIF D4 (Šídák post-hoc) | DF = 234 | > 0.9999 |
| | | Females: Vehicle v MIF D5 (Šídák post-hoc) | DF = 234 | > 0.9999 |
| | | Females: Vehicle v MIF D6 (Šídák post-hoc) | DF = 234 | > 0.9999 |
| | | Females: Vehicle v MIF D7 (Šídák post-hoc) | DF = 234 | > 0.9999 |
| | | Females: Vehicle v MIF D8 (Šídák post-hoc) | DF = 234 | > 0.9999 |

| | | | | |
|---|---|---|---|---|
| **Fig. S4D** | Lever pressing rate | Training Day | F(8, 208) = 67.32 | < 0.0001 |
| | | Sex | F(1, 26) = 18.38 | 0.0002 |
| | | Treatment | F(1, 26) = 7.004 | 0.0136 |
| | | Training day x Sex | F(8, 208) = 8.074 | < 0.0001 |
| | | Training Day x Treatment | F(8, 208) = 4.769 | < 0.0001 |
| | | Sex x Treatment | F(1, 26) = 16.06 | 0.0005 |
| | | Training Day x Sex x Treatment | F(8, 208) = 11.88 | < 0.0001 |
| **Fig. S4D** | Lever pressing rate | Males: Vehicle v MIF D0 (Šídák post-hoc) | DF = 234 | > 0.9999 |
| | | Males: Vehicle v MIF D1 (Šídák post-hoc) | DF = 234 | > 0.9999 |
| | | Males: Vehicle v MIF D2 (Šídák post-hoc) | DF = 234 | > 0.9999 |
| | | Males: Vehicle v MIF D3 (Šídák post-hoc) | DF = 234 | > 0.9999 |
| | | Males: Vehicle v MIF D4 (Šídák post-hoc) | DF = 234 | > 0.9999 |
| | | Males: Vehicle v MIF D5 (Šídák post-hoc) | DF = 234 | 0.0187 |
| | | Males: Vehicle v MIF D6 (Šídák post-hoc) | DF = 234 | < 0.0001 |
| | | Males: Vehicle v MIF D7 (Šídák post-hoc) | DF = 234 | < 0.0001 |
| | | Males: Vehicle v MIF D8 (Šídák post-hoc) | DF = 234 | < 0.0001 |
| **Fig. S4D** | Lever pressing rate | Females: Vehicle v MIF D0 (Šídák post-hoc) | DF = 234 | > 0.9999 |
| | | Females: Vehicle v MIF D1 (Šídák post-hoc) | DF = 234 | > 0.9999 |
| | | Females: Vehicle v MIF D2 (Šídák post-hoc) | DF = 234 | > 0.9999 |
| | | Females: Vehicle v MIF D3 (Šídák post-hoc) | DF = 234 | > 0.9999 |
| | | Females: Vehicle v MIF D4 (Šídák post-hoc) | DF = 234 | > 0.9999 |
| | | Females: Vehicle v MIF D5 (Šídák post-hoc) | DF = 234 | > 0.9999 |

| | | | | |
|---|---|---|---|---|
| | | Females: Vehicle v MIF D6 (Šídák post-hoc) | DF = 234 | > 0.9999 |
| | | Females: Vehicle v MIF D7 (Šídák post-hoc) | DF = 234 | > 0.9999 |
| | | Females: Vehicle v MIF D8 (Šídák post-hoc) | DF = 234 | 0.9384 |
| **Fig. S4E** | Training session acquisition | Training Session | F (2, 78) = 18.14 | < 0.0001 |
| | | Sex | F (1, 78) = 0.7299 | 0.3955 |
| | | Treatment | F (1, 78) = 0.02273 | 0.8806 |
| | | Training Session x Sex | F (2, 78) = 0.5537 | 0.5771 |
| | | Training Session x Treatment | F (2, 78) = 0.005682 | 0.9943 |
| | | Sex x Treatment | F (1, 78) = 1.578 | 0.2127 |
| | | Training Session x Sex x Treatment | F (2, 78) = 1.311 | 0.2753 |
| **Fig. S4F, S4G** | Test press rates Males and Females | Sex | F(2, 35) = 25.48 | 0.0446 |
| | | Treatment | F(1, 35) = 4.3342 | < 0.0001 |
| | | Sex x Treatment | F(2, 35) = 0.5438 | 0.5854 |
| **Fig. S4F** | Male extinction test | Veh_Male v CORT_Male (Tukey post-hoc) | DF = 35 | < 0.0001 |
| | | Veh_Male v CORT+MIF_Male (Tukey post-hoc) | DF = 35 | 0.8193 |
| | | CORT_Male v CORT+MIF_Male (Tukey post-hoc) | DF = 35 | 0.0006 |
| **Fig. S4G** | Female extinction test | Veh_Female v CORT_Female (Tukey post-hoc) | DF = 35 | 0.0054 |
| | | Veh_Female v CORT+MIF_Female (Tukey post-hoc) | DF = 35 | 0.9991 |
| | | CORT_Female v CORT+MIF_Female (Tukey post-hoc) | DF = 35 | 0.007 |
| **Fig. S4H** | Consumption test | Pre v Post | F(1, 26) = 224.9 | < 0.0001 |

| | | | | |
|---|---|---|---|---|
| | | Sex | F(1, 26) = 12.87 | 0.0014 |
| | | Treatment | F(1, 26) = 0.3385 | 0.5657 |
| | | Pre v Post x Sex | F(1, 26) = 4.289 | 0.484 |
| | | Pre v Post x Treatment | F(1, 26) = 0.4027 | 0.5312 |
| | | Sex x Treatment | F(1, 26) = 4.974 | 0.0346 |
| | | Pre v Post x Sex x Treatment | F(1, 26) = 4.57 | 0.0421 |

**Table S4: Detailed statistics for Figure S4.** Either two-way or three-way analysis of the variance (ANOVA) tests were run with either a Šidák or Tukey post-hoc to correct for multiple comparisons. Corresponding figures, metrics, ANOVA comparisons, F statistics, and p values are shown.

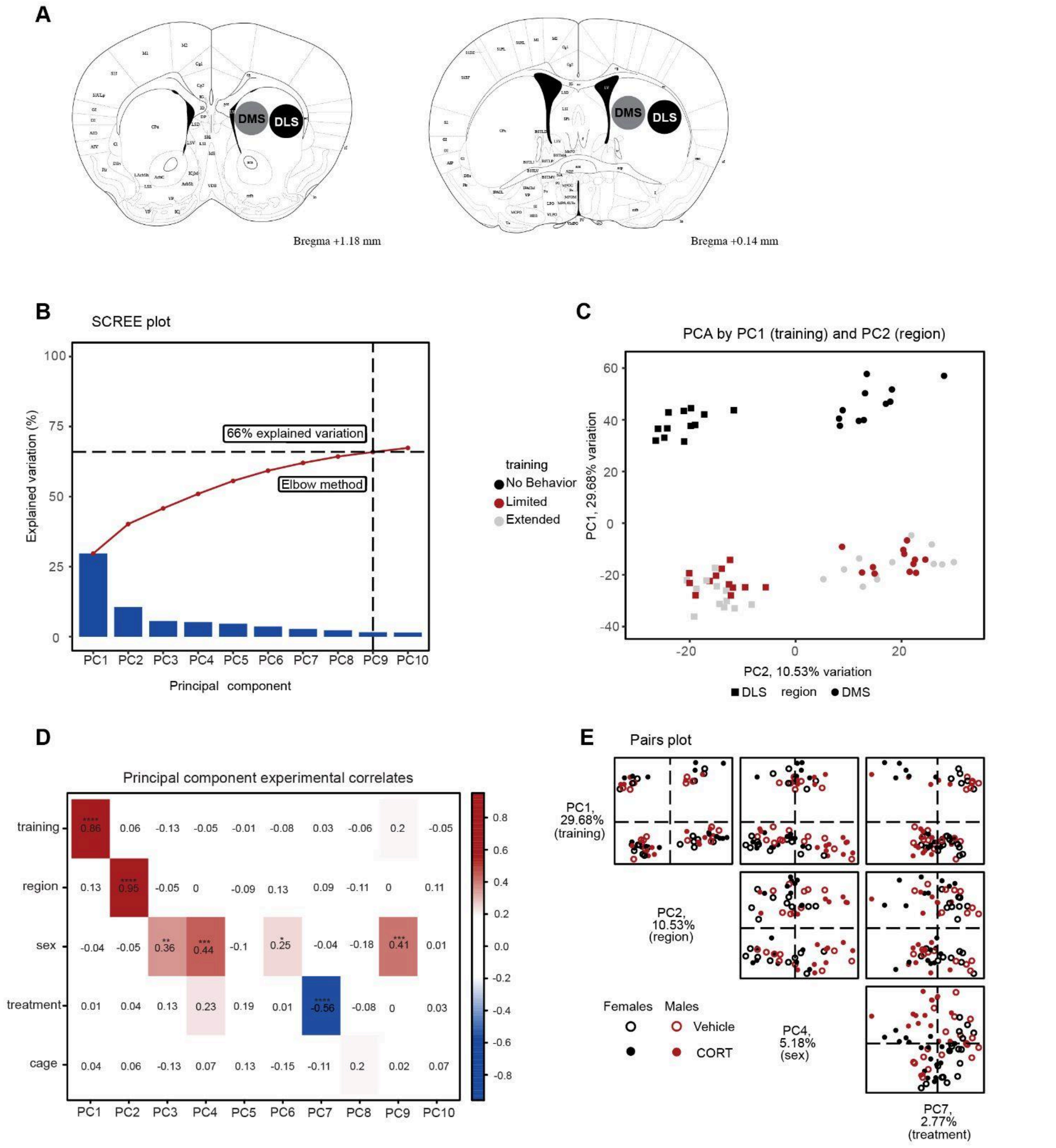

**Figure S5**: **Behavioral timeline with tissue recovery locations and principal component analyses of RNA sequencing.** **A** Coronal slices from the Paxinos Mouse Brain Atlas [11] showing tissue recovery locations. The left hemisphere has known anatomical structures, while the right hemisphere highlights the locations selected for the 2mm tissue punch biopsies of the DMS and DLS. **B** A scree plot displaying the amount of experimental variation explained by each principal component. The elbow method was used to find the inflection point (PC9) at which all additional principal components add an insignificant amount of variation. Up to 66% of experimental variation is explained at this inflection point. **C** A biplot of individual samples organized in space by the top two principal components. PC1 splits samples by training (no training vs any amount of training (Limited + Extended)), while PC2 splits samples by region (DLS or DMS). **D** A Pearson's correlation between principal components and known experimental correlates. Training (PC1), region (PC2), sex (PC3, PC4, PC6, PC9), and treatment (PC7) significantly correlate with principal components, while no significant cage/cohort difference can be detected by a principal component analysis. **E** A pairs plot displaying the most significant principal components by experimental correlate is plotted in pairs to show individual subject segregation in space by each combination of principal components. Training (PC1), brain region (PC2), sex (PC4), and CORT treatment (PC7) are shown in axis pairs along with the relative percentage of variability explained for each PC.

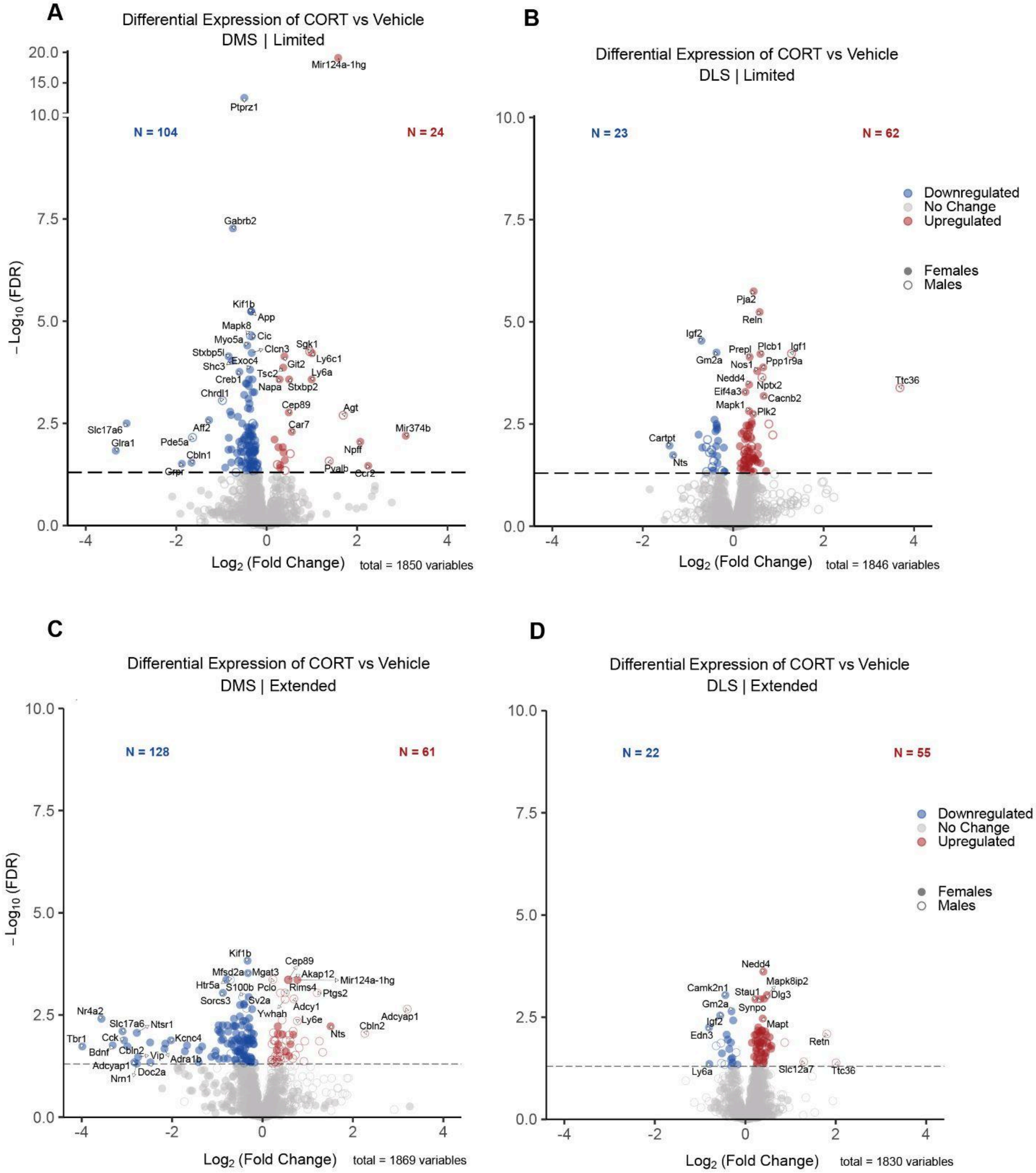

**Figure S6: Differential gene expression from CORT versus vehicle groups following limited training.** **A–B** Expanded Figure 2C. **A–D** Differentially expressed genes (DEGs) across 1157 genes from gene ontology terms: "synaptic signaling" and "cognition." Significance cutoff (horizontal line at FDR = 0.05). Differential gene expression from CORT versus vehicle groups produces a decrease in plasticity gene expression in the DMS and an increase in plasticity gene expression in the DLS.

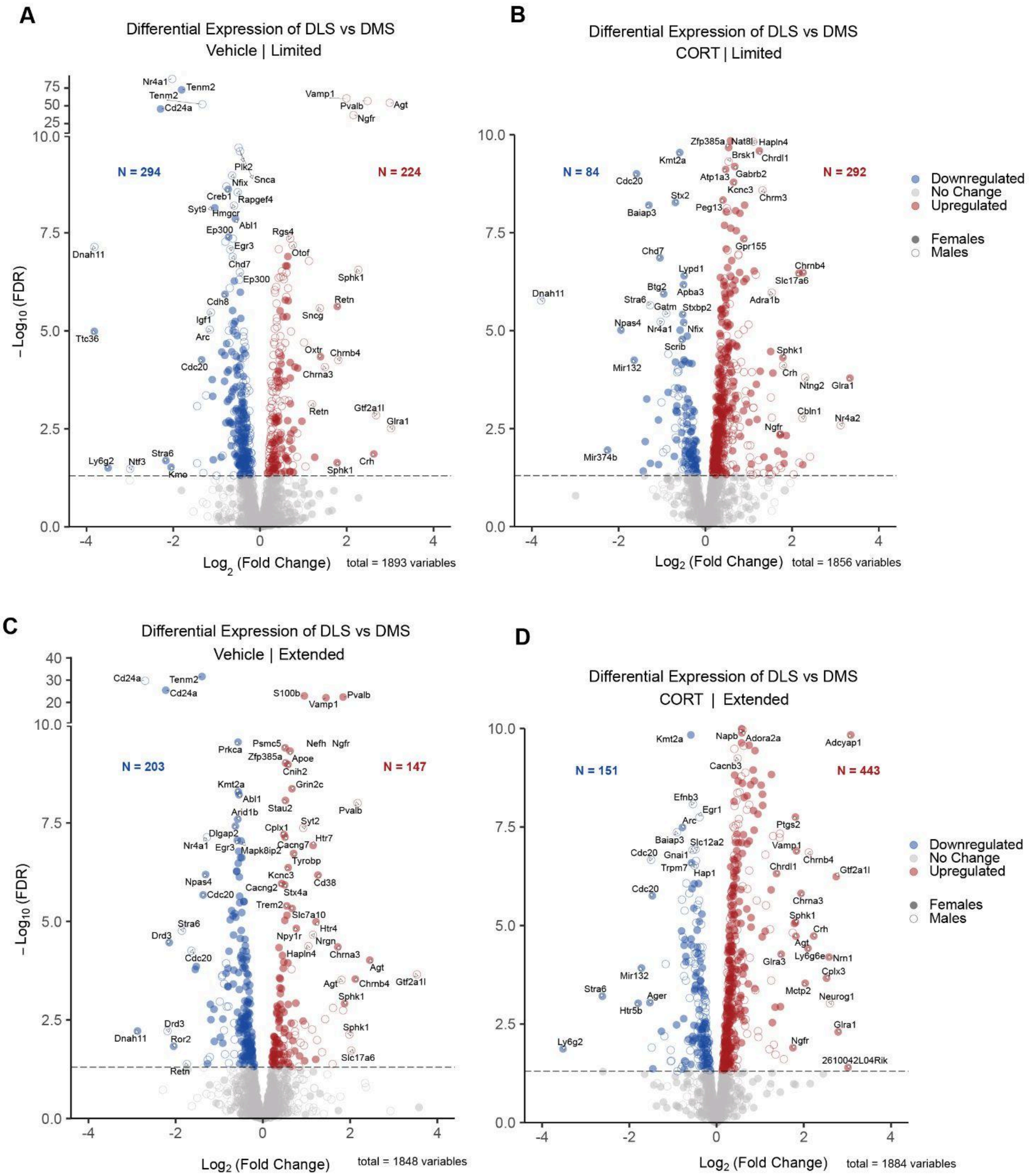

**Figure S7: Differential gene expression from DLS versus DMS groups following limited training. A–B** Expanded Figure 2D. **A–D** Differentially expressed genes (DEGs) across 1157 genes from gene ontology terms: “synaptic signaling” and “cognition.” Significance cutoff (horizontal line at FDR = 0.05). Differential gene expression from DLS versus DMS groups produces similar amounts of up and downregulated genes in vehicle groups and produces less downregulation and more upregulation in CORT-treated groups.

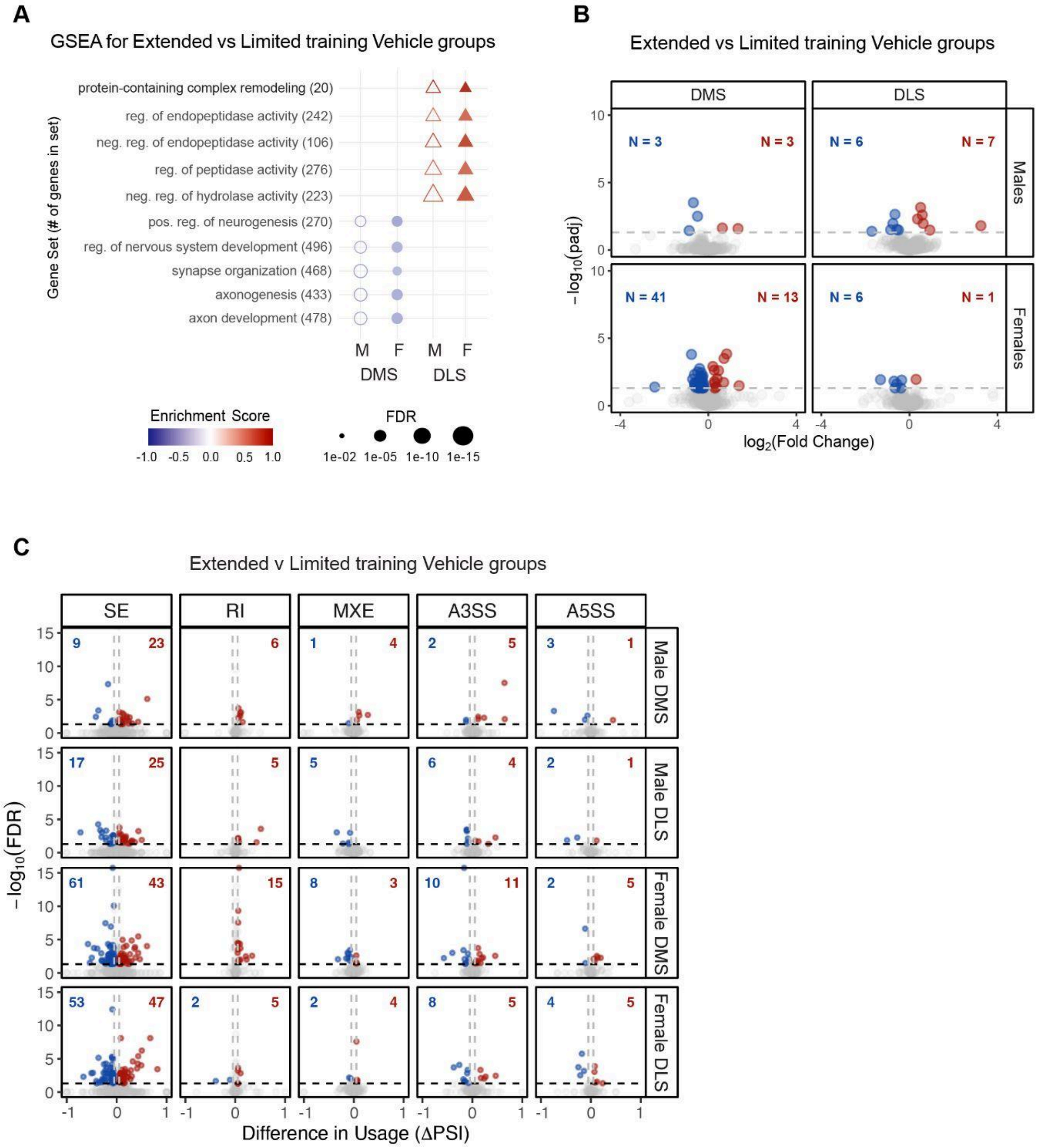

**Figure S8: The dorsal striatum undergoes sex- and subregion-specific gene regulation as behaviors become routinized.** **A** Gene set enrichment analysis for extended versus limited training based on differential gene expression. Term selection was restricted to the five lowest FDRs shared among male and female DMS or DLS. The DMS had a deenrichment of synaptic plasticity-related terms in both male and female mice following extended training compared to limited training. The DLS had an enrichment of activity-regulation-related terms. **B** Differential gene expression of extended training compared to limited training groups by region and sex (n = 3/training duration/sex/region). Differentially expressed genes (DEGs) across 1157 genes from gene ontology terms: “synaptic signaling” and “cognition.” Significance cutoffs (horizontal line at

FDR = 0.05, vertical lines at |ΔPSI| > 5%). Vehicle male DMS, male DLS, and female DLS exhibited low amounts of differential plasticity gene expression. The female DMS exhibited the most amount of differential plasticity gene expression, most of which was downregulated. **C** Alternative splice events across 1157 genes from gene ontology terms: "synaptic signaling" and "cognition." Significance cutoff (horizontal line at FDR = 0.05). Extended training, when compared to limited training, produced increased and decreased inclusion of spliced exons (SEs) in the DMS and DLS of male and female mice, and increased the inclusion of retained introns (RIs) in the DMS and DLS of male and female mice. There was dysregulation but no established trend for differences in mutually exclusive exons (MXE), alternative 3' splice site (A3SS), or alternative 5' splice site (A5SS).

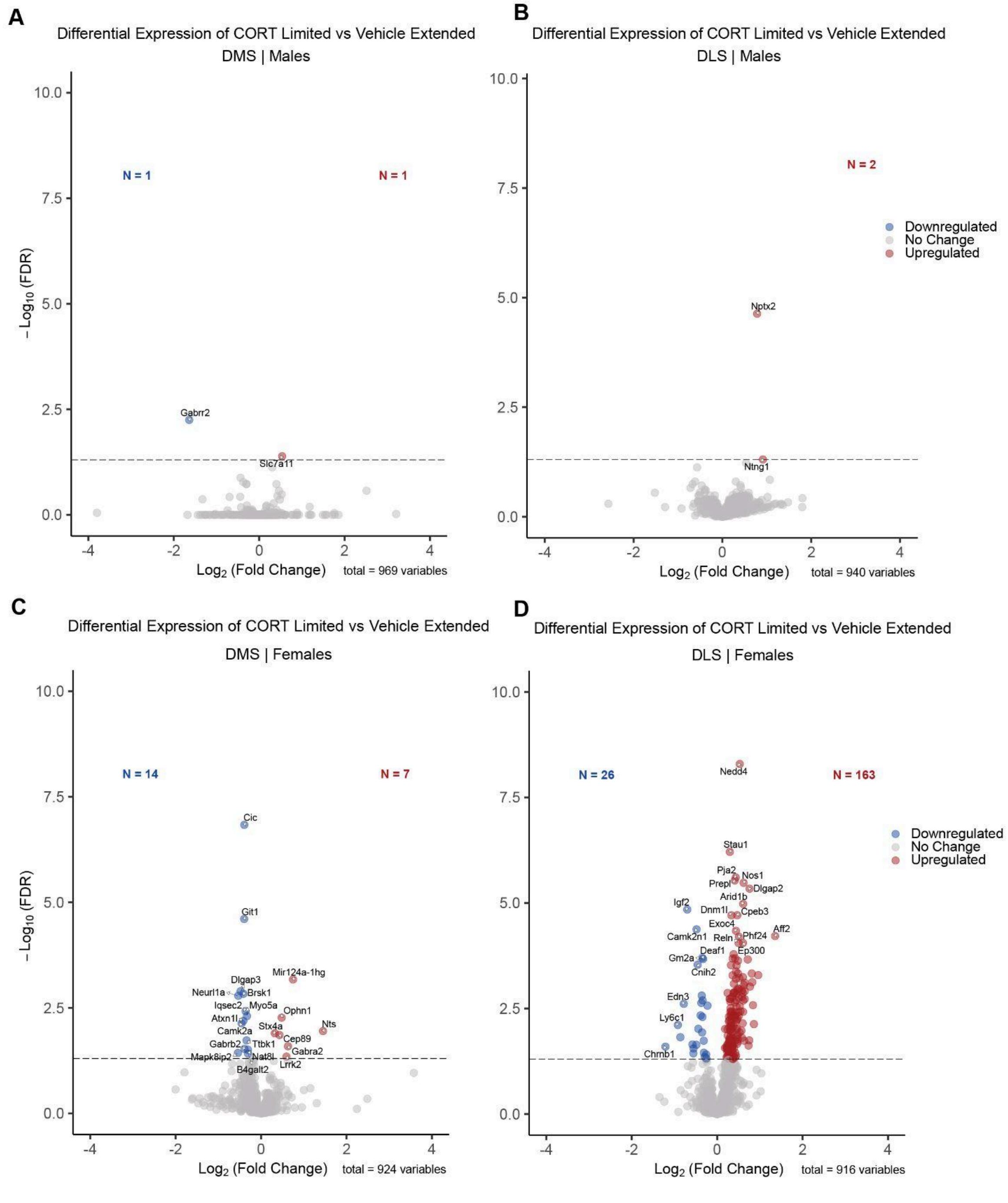

**Figure S9: Differential gene expression from CORT-limited training versus vehicle-extended training groups. A–D** Expanded Figure 2E. Differentially expressed genes (DEGs) across 1157 genes from gene ontology terms: “synaptic signaling” and “cognition.” Significance cutoff (horizontal line at FDR = 0.05). Differential gene expression from CORT-limited versus vehicle-extended groups produces low differences in male DMS and DLS plasticity gene expression. Meanwhile, the same comparison produces some differences in female DMS and substantial differences in female DLS plasticity gene expression.

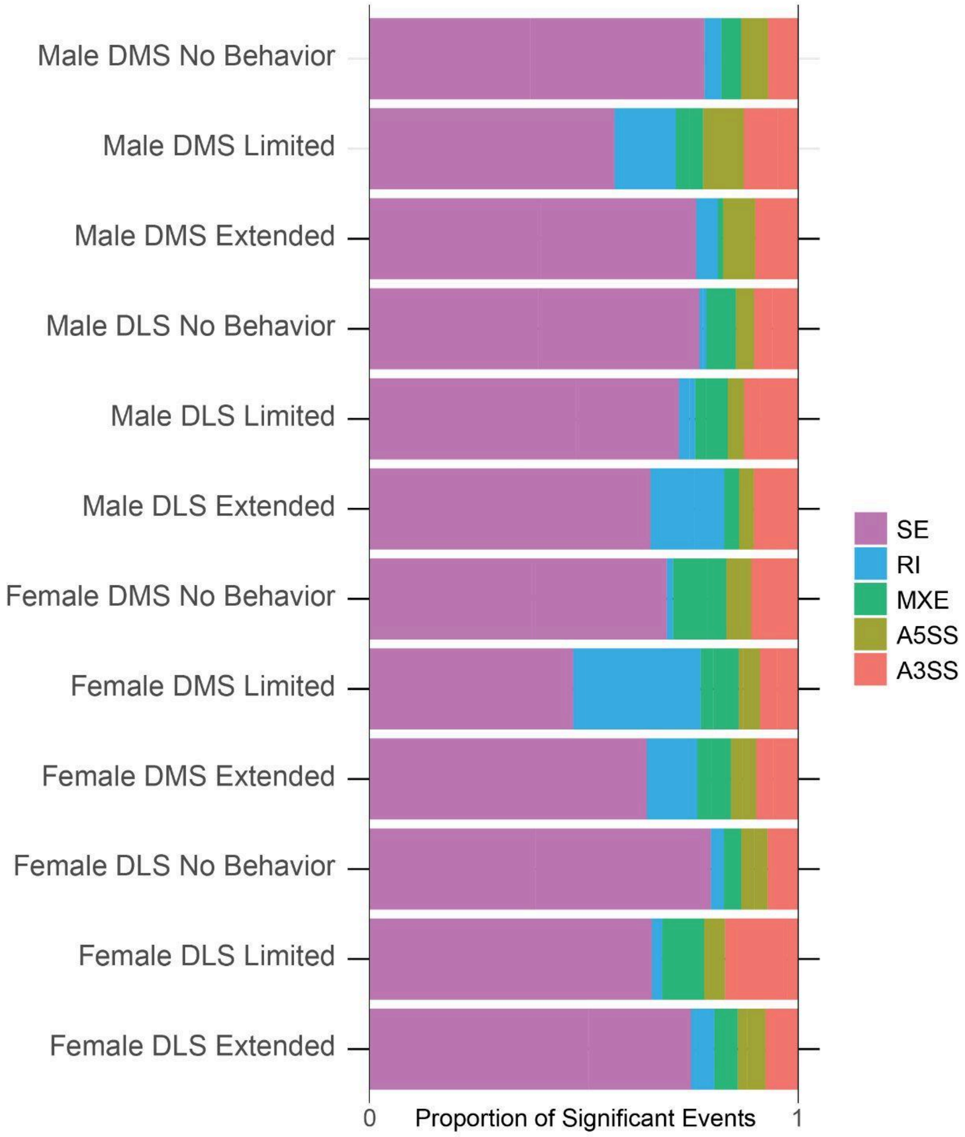

**Figure S10: CORT treatment produces differential alternative splicing across sex, region, and training time point.** Relative proportions of differential splice events from CORT versus vehicle comparisons across sex, region, and training time point (n = 3/sex/region/treatment/training). Alternative splicing events: spliced exon, SE, retained intron, RI, mutually exclusive exon, MXE, alternative 3' splice site, A3SS, or alternative 5' splice site, A5SS. SEs were the most abundant differential splicing event due to CORT treatment across all groups, followed by RIs and A3SSs.

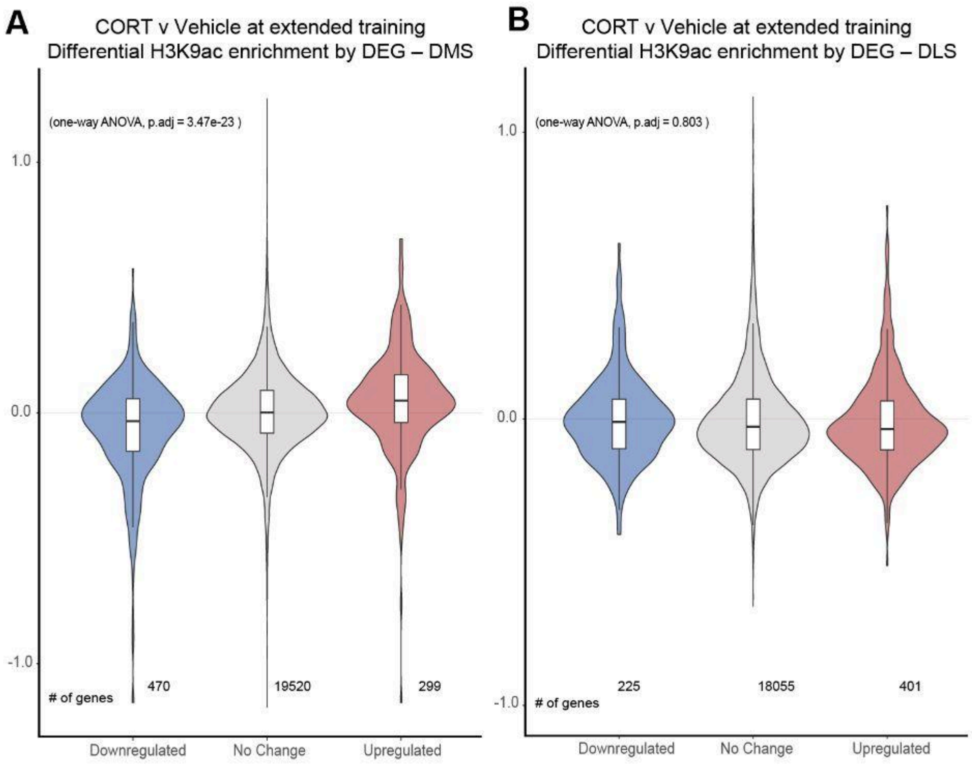

**Figure S11: CORT treatment produces differential H3K9ac enrichment related to differentially expressed genes by region during extended training.** Differential H3K9ac enrichment (n = 3/sex/region/treatment/training) and DEGs (n = 3/sex/region/treatment/training) were calculated by CORT versus vehicle comparisons. The log2 fold change of differential H3K9ac enrichment is plotted by direction of change for the DEGs. **A, B** CORT led to a significant correlation between H3K9ac and DEGs in DMS, but not DLS, following extended training.

| Figure | Metric | ANOVA comparison | F(DFn, DFd) | p value |
|---|---|---|---|---|
| **Fig. 4C** | H3K9ac enrichment related to DEGs – Limited training DMS | DEG | F(2, 19745) = 16.535 | 6.68E-08 |
| **Fig. 4D** | H3K9ac enrichment related to DEGs – Limited training DLS | DEG | F(2, 11621) = 1.095 | 0.335 |
| **Fig. S11A** | H3K9ac enrichment related to DEGs – Extended training DMS | DEG | F(2, 20286) = 51.849 | 3.47E-23 |
| **Fig. S11B** | H3K9ac enrichment related to DEGs – Extended training DLS | DEG | F(2, 18678) = 0.219 | 0.803 |

**Table S5: Detailed statistics for Figures 4C–D and Figure S11.** One-way analysis of the variance (ANOVA) tests were run for all genes aggregated by differentially expressed gene (DEG) designation (Downregulated, No Change, Upregulated). Corresponding figures, metrics, ANOVA comparisons, F statistics, and p values are shown.

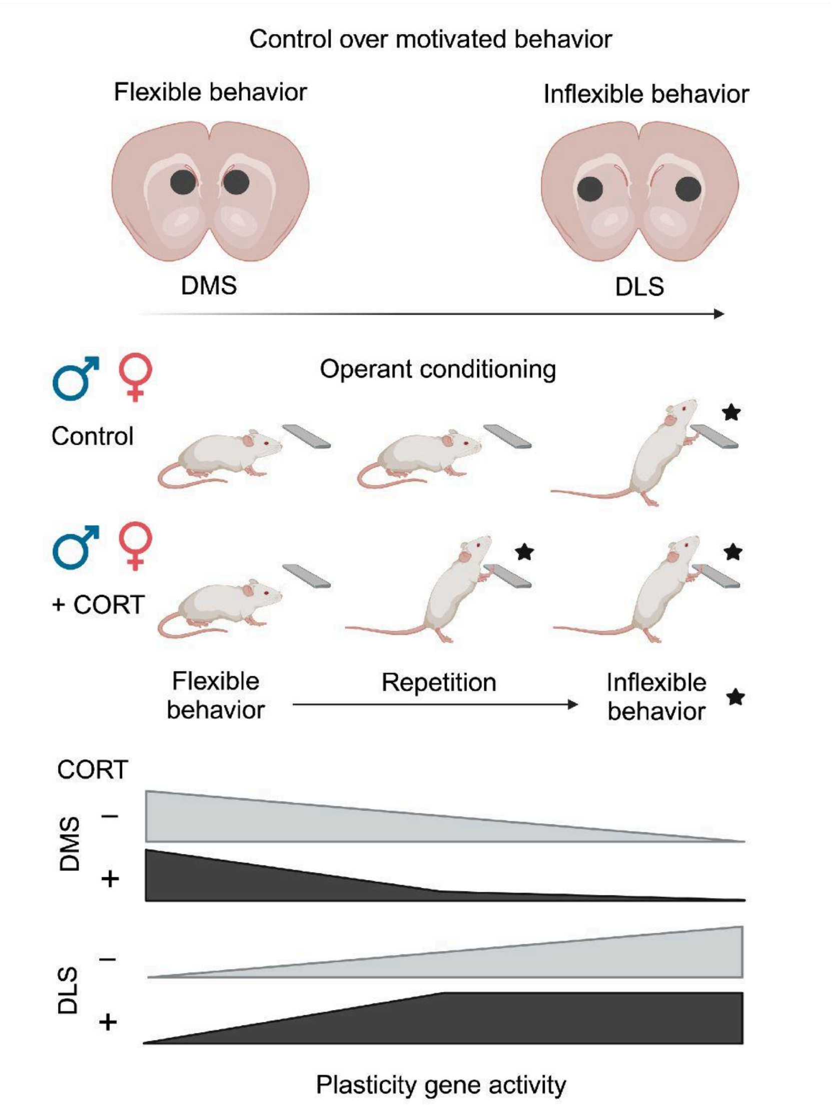

**Figure S12**: **Proposed graphical abstract of CORT's influence on behavioral inflexibility and plasticity gene activity.** Control over motivated behavior is canonically distributed between the DMS and DLS. The DMS promotes behavioral flexibility, and the DLS promotes behavioral inflexibility. Increasing the number of operant learning sessions increases DLS-dependent behavioral inflexibility. Chronic CORT treatment hastens the transition to behavioral inflexibility in rodents. Plasticity gene regulation is also affected by chronic CORT, in a subregion-specific manner, during limited amounts of operant training. Chronic CORT reduces plasticity gene activity in the DMS and increases the plasticity gene activity in the DLS during limited training, likely contributing to behavioral inflexibility.
